# Netrin-1 binding to Unc5B regulates Blood-Retina Barrier integrity

**DOI:** 10.1101/2023.01.21.525006

**Authors:** Jessica Furtado, Luiz Henrique Geraldo, Felipe Saceanu Leser, Mathilde Poulet, Hyojin Park, Laurence Pibouin-Fragner, Anne Eichmann, Kevin Boyé

## Abstract

**Background:** The blood brain barrier (BBB) preserves neuronal function in the central nervous system (CNS) by tightly controlling metabolite exchanges with the blood. In the eye, the retina is likewise protected by the blood-retina barrier (BRB) to maintain phototransduction. We showed that the secreted guidance cue Netrin-1 regulated BBB integrity, by binding to endothelial Unc5B and regulating canonical β-catenin dependent expression of BBB gene expression

**Objective:** Here, we investigated if Netrin-1-binding to endothelial Unc5B also controlled BRB integrity, and if this process involved Norrin/β-catenin signaling, which is the major known driver of BRB development and maintenance.

**Methods:** We analyzed Tamoxifen-inducible loss- and gain-of-function alleles of *Unc5B, Ntn1* and *Ctnnb1* in conjunction with tracer injections and biochemical signaling studies.

**Results:** Inducible endothelial *Unc5B* deletion, and inducible global *Ntn1* deletion in postnatal mice reduced phosphorylation of the Norrin receptor LRP5, leading to reduced β-catenin and LEF1 expression, conversion of retina endothelial cells from a barrier-competent Claudin-5+/PLVAP-state to a Claudin-5-/PLVAP+ leaky phenotype, and extravasation of injected low molecular weight tracers. Inducible *Ctnnb1* gain of function rescued vascular leak in *Unc5B* mutants, and *Ntn1* overexpression induced BRB tightening. Unc5B expression in pericytes contributed to BRB permeability, via regulation of endothelial Unc5B. Mechanistically, Netrin-1-Unc5B signaling promoted β-catenin dependent BRB signaling by enhancing phosphorylation of the Norrin receptor LRP5 via the Discs large homologue 1 (Dlg1) intracellular scaffolding protein.

**Conclusions:** The data identify Netrin1-Unc5B as novel regulators of BRB integrity, with implications for diseases associated with BRB disruption.

## INTRODUCTION

Blood vessels in the central nervous system (CNS) form a specialized blood-brain barrier (BBB) that prevents entry of toxins and tightly controls the movement of ions, molecules, and cells between the blood and the brain, thereby ensuring neuronal function and protection^1-5^. As part of the CNS, the light-sensitive retina at the back of the eye is likewise protected by the inner blood-retina barrier (BRB). The impermeability of the BBB and BRB impedes drug and treatment delivery during various pathologies^6-9^. Conversely, sustained loss of barrier integrity leads to the development of multiple diseases including stroke, diabetic retinopathy and retinal inflammatory disease^4,5,10-12^. Overall, the development of tailored therapeutic strategies designed to increase or reduce CNS barrier permeability “on-demand” is a critical yet unmet clinical need.

Canonical Wnt/Norrin/β-catenin signaling is the main known pathway that maintains CNS ECs in barrier-competent state^13-16^. The specific Wnt/Norrin ligands and receptors that maintain barrier properties differ in a region-specific manner within the CNS. Cerebral cortex ECs utilize Wnt7a/7b signaling through LRP6, Frizzled4, and two Wnt7a/7b specific cell-surface co-activators GPR124 and RECK^13,14,17-21^. By contrast, retina and cerebellar ECs rely on Norrin signaling through LRP5, Frizzled4 and the Norrin cell-surface co-activator TSPAN12^13,14,17,18,22,23^. Receptor activation converges on β-catenin stabilization and nuclear translocation to induce activation of TCF and LEF1 transcription factors, thereby regulating expression of a barrier-specific gene transcription repertoire, including the tight junction associated protein Claudin-5 that suppresses paracellular permeability^24,25^, influx and efflux transporters that control exchange between the blood and the tissue^26-28^, while inhibiting expression of the permeability protein PLVAP that forms EC fenestrae and transcytotic vesicles^13,14,29^. Signaling downstream of Norrin and Wnt7a/7b depend on the Discs large homologue 1 (Dlg1) intracellular scaffolding protein, which enhances both Wnt-mediated and Norrin-mediated β-catenin stabilization and signaling at the BBB and BRB through as yet unclear mechanisms^30^.

Netrin-1 is a laminin-related secreted protein initially discovered for guidance of commissural axons during embryonic development^31^ that is emerging as a multifunctional cue in diverse biological processes including vascular development^32-34^ inflammation^35-40^and neuroprotection^41,42^. Netrin-1 modulates inflammation in atherosclerosis^36-38^ and obesity^40^. In mouse models of multiple sclerosis, Netrin-1 treatment was shown to be neuroprotective by stabilizing the BBB^39^, and Netrin-1 also induced survival of various neuronal populations, including dopaminergic neurons^43^.

Netrin-1 signals to inflammatory cells, glia, neurons and endothelial cells via several transmembrane receptors: Deleted in colorectal cancer (Dcc)^44^, Neogenin, A2B^45^ and Unc5^46^, which are expressed by different cell types in the adult brain including ECs (Unc5B), oligodendrocytes (Unc5B, Neogenin and A2B) and astrocytes (Neogenin, A2B)^47^. Among those, Unc5B is predominantly expressed in endothelial cells in mice and humans^34^. Besides Netrin-1, Unc5b also binds to endothelial Robo4^48^ and Flrt2^49^. The existence of several Netrin-1 receptors, and various Unc5B ligands, has rendered understanding of their signaling interactions in the adult and in pathologies difficult, a problem that is compounded by the lethality of global ko mice^31,34^. To tackle this issue, we reported generation of inducible *Ntn1* and *Unc5B* mutants^50,51^, and we observed that adult deletion of *Ntn1* and of endothelial *Unc5B* produced a BBB leak that was characterized by reduced brain EC LRP6 phosphorylation and β-catenin signaling and rescued by genetic β-catenin overexpression^51^, identifying Netrin-1 signaling via Unc5B as a novel mediator of BBB integrity signaling that cross talks with canonical Wnt signaling.

Herein, we investigated if Unc5B mediated CNS barrier protection extended to the BRB and required Netrin-1 and Norrin signaling through LRP5 and β-catenin signaling.

## RESULTS

### Unc5B is expressed in endothelial cells and mural cells in the postnatal retina

Retinal angiogenesis in mice begins after birth and is complete after three weeks^52^. At postnatal day (P)1 the superficial vascular plexus starts to sprout from the optic nerve, and it reaches the edges of the retina at around P7. Tip cells guiding this superficial sprouting are deprived of a functional BRB to allow proper guidance and vascular development and barrier properties are gradually formed as vessels mature^53^. At P7, the Norrin/β-catenin signaling pathway induces the formation of diving tip cells that sprout vertically to form the deep and the intermediate vascular plexus^22,54^. These diving vessels form coincident with the onset of vision and possess a functional BRB^53,55^.

Single-cell RNA sequencing data from retinas collected at P6 and P10^55^ showed that *Unc5B* was expressed at highest levels by ECs and mural cells at both stages (**Fig. 1a,b and Supp. Fig. 1a**). Among the EC cluster, *Unc5B* was expressed in a zonated manner, with arterial ECs expressing highest levels, followed by capillaries that expressed lower *Unc5B* levels, and veins that exhibited essentially absent *Unc5B* mRNA (**Fig 1c,d and Supp. Fig. 1b**). *Unc5B* was also strongly expressed by tip cells in both the superficial vascular layer at P6, and in diving D tip cells that invade the neuroretina at P10 (**Fig 1c,d and Supp. Fig. 1b**). Whole-mount retina immunostaining at P5 and P12 confirmed Unc5B protein expression by arterial ECs, followed by tip cells and capillaries but was not detected in veins (**Fig. 1e-g**), Higher magnification images also revealed Unc5B expression in CD13+pericytes at somewhat lower levels when compared with ECs (**Fig. 1h**). The Unc5B ligand Flrt2 was expressed by retinal pigment epithelium (RPE) and venous and capillary ECs, while Robo4 was specifically expressed by ECs (**Supp. Fig. 1c**). Interestingly, retina cells exhibited essentially absent *Netrin-1* mRNA (**Supp. Fig. 1c**).

**Figure 1:**
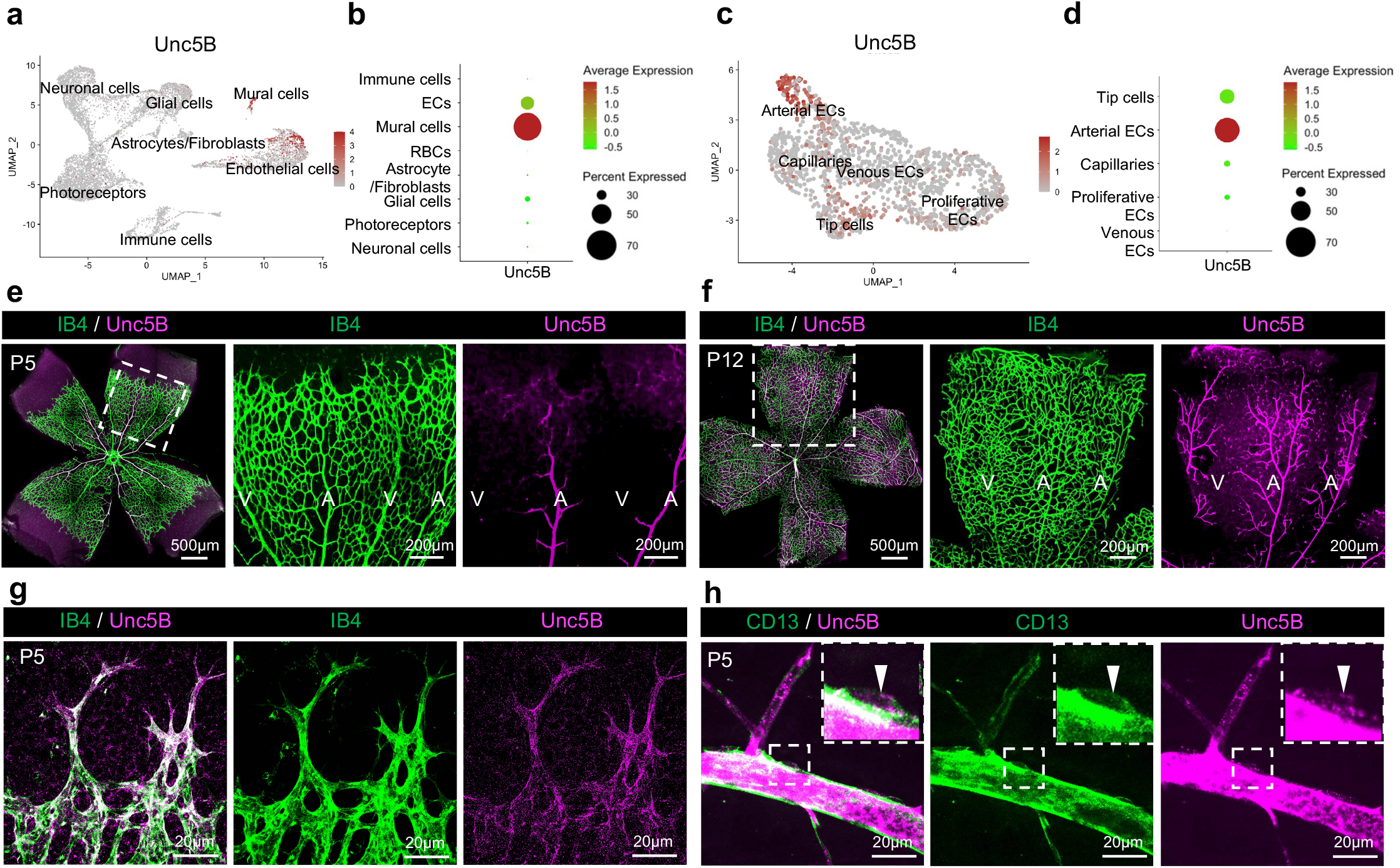
Unc5B expression in endothelial cells and mural cells of the developing retina. (a) UMAP plot of retinal cells^55^. (b) Unc5B dot plot expression levels and frequency among retina cell clusters. Color scale: red: high expression; green: low expression. (c) UMAP plot of EC sub-clusters. (d) Unc5B dot plot expression levels and frequency among ECs subclusters. Color scale: red: high expression; green: low expression. (e-h) Immunofluorescence staining and confocal imaging on whole-mount P5 or P12 retinas with the indicated antibodies.

### Endothelial Unc5B regulates retinal angiogenesis and BRB formation

To investigate Unc5B function in retinal ECs, we used tamoxifen (TAM)-inducible endothelial-specific *Unc5B* knockout mice (hereafter *Unc5BiECko*) that were generated by crossing *Unc5B*^*fl/fl*^ mice^51^ with *CDH5Cre*^*ERT2*^ mice^56^. Gene deletion was induced by injecting TAM from P0-2 and immunostaining revealed endothelial-specific deletion in *Unc5BiECko* mice (**Supp. Fig. 2a-c**). Endothelial *Unc5B* deletion induced transient vascular development defects, with reduced vascular outgrowth and increased vascular density at P5 compared to TAM-injected cre-negative littermate controls (**Fig. 2a-c**). When we induced gene deletion by TAM injection at P6-8 and sacrificed pups at P12, *Unc5BiECko* mice showed superficial, intermediate, and deep plexus densities that were similar to TAM-treated cre-negative littermate controls (**Fig. 2d-f**). Hence, Unc5B transiently controls retina vascular development.

**Figure 2:**
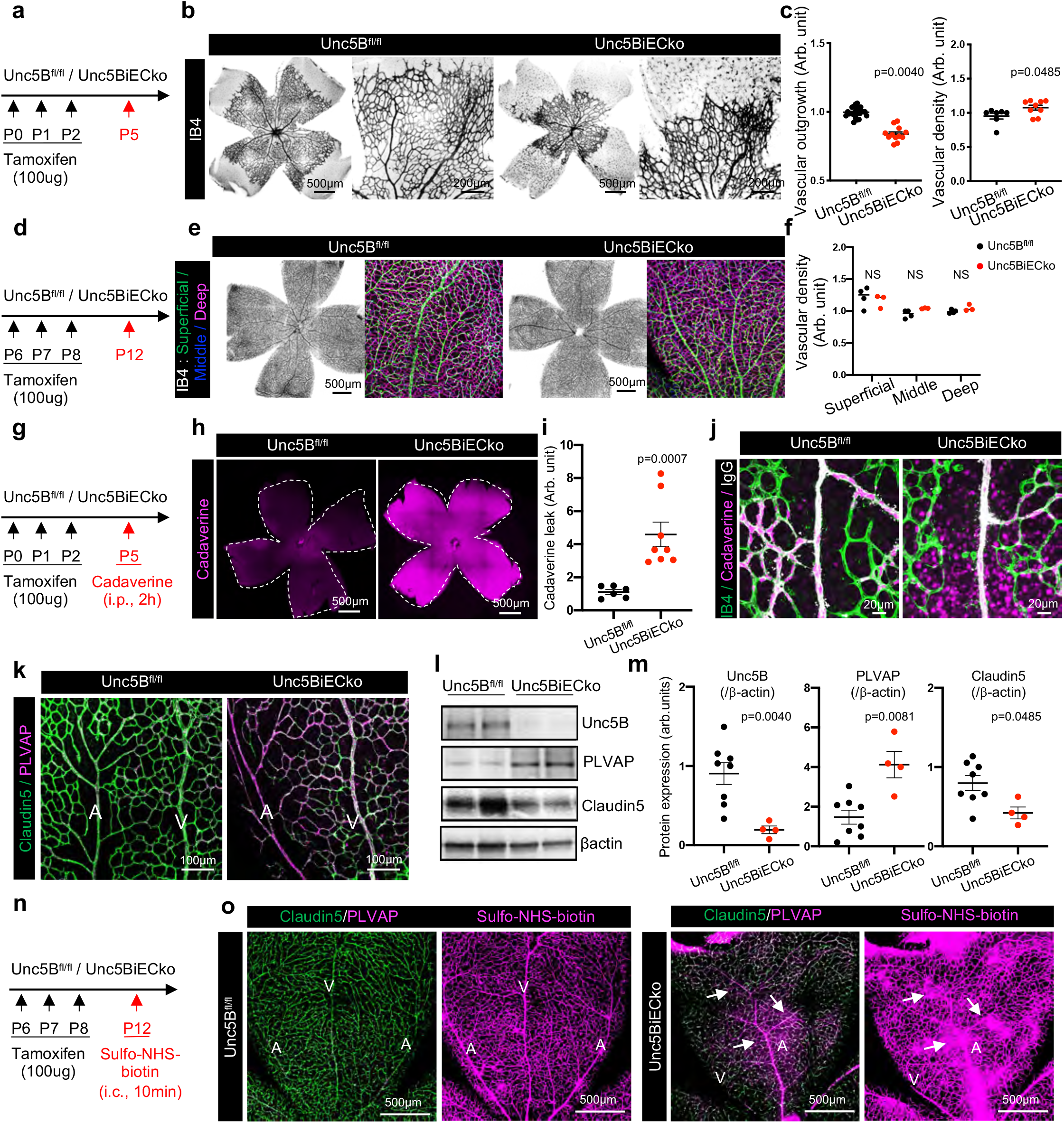
Endothelial Unc5B regulates retinal angiogenesis and BRB formation. (a) *Unc5B* gene deletion strategy using tamoxifen injection in P5 neonates. (b) IB4 immunofluorescence staining and confocal imaging on whole-mount P5 retinas. (c) Quantification of IB4+ vascular outgrowth and density in P5 retinas. (d) *Unc5B* gene deletion strategy using tamoxifen injection in P12 neonates. (e) Immunofluorescence staining of IB4+ vessel and confocal imaging on whole-mount P12 retinas. Superficial, intermediate and deep plexus are color-coded. (f) Quantification of superficial, intermediate and deep plexus vascular density in P12 retinas. (g) *Unc5B* gene deletion strategy using tamoxifen injection in P5 neonates. (h) Confocal imaging on whole-mount P5 retinas, 2h after i.p. cadaverine injection. (i) quantification of cadaverine leakage. (j) Immunofluorescence staining and confocal imaging on whole-mount P5 retinas with the indicated antibodies, 2h after i.p. cadaverine injection. (k) Immunofluorescence staining and confocal imaging on whole-mount P5 retinas with the indicated antibodies. (l,m) Western blot (l) and quantification (m) of P5 retina protein extracts. (n) *Unc5B* gene deletion strategy using tamoxifen injection in P12 neonates. (o) Immunofluorescence staining and confocal imaging on whole-mount P12 retinas with the indicated antibodies, 10min after i.v. sulfo-NHS-biotin injection. All data are shown as mean+/-SEM. Two-sided Mann-Whitney U test was performed for statistical analysis.

As we previously showed that TAM treatment between P0-2 led to BBB disruption, seizures, and lethality 10-12 days after gene deletion^51^, we sought to determine if Unc5B also regulates BRB permeability. We induced gene deletion at P0-2 and probed BRB integrity at P5 by intraperitoneal (i.p.) cadaverine injection for 2h followed by PBS cardiac perfusion and retina whole-mount (**Fig. 2g**). Cadaverine leakage was observed in *Unc5BiECko* retinas compared to TAM-treated cre-negative littermate controls (**Fig. 2h,i**), indicating that endothelial Unc5B is a positive regulator of BRB integrity. Injection of cadaverine, followed by 150kDa mouse IgG immunostaining revealed a size-selective opening of the BRB upon endothelial Unc5B deletion as cadaverine, but not mouse IgG leaked through the BRB (**Fig. 2j**).

To determine the cause of the BRB defect, we examined expression levels and localization of proteins relevant for BRB function in ECs, pericytes and astrocytes. Compared to TAM-treated cre-negative littermate controls, *Unc5BiECko* retinas had similar PDGFRβ and GFAP expression pattern, attesting similar pericyte and astrocyte coverage, respectively (**Supp. Fig. 2d-f**). By contrast, *Unc5BiECko* P5 retinas exhibited significantly reduced expression of the tight-junction protein Claudin-5, along with increased expression of the permeability protein PLVAP in arteries and capillaries (**Fig. 2k**), suggesting that BRB leakage may be due to changes in expression of Claudin-5 and PLVAP. Western blotting of mouse retina protein lysates confirmed decreased Claudin-5 and increased PLVAP protein expression in *Unc5BiECko* P5 neonates compared to cre-negative littermate controls (**Fig. 2l,m**).

We sought to check if Unc5B-mediated BRB defects are lasting, by inducing gene deletion in mice at P6,7 and 8 followed by i.v. injection of sulfo-NHS-biotin tracer and retina whole-mount at P12 (**Fig. 2n**). Sulfo-NHS-biotin remained inside the Claudin-5+/PLVAP-BRB-competent vasculature of TAM-injected cre-negative littermate controls but leaked in *Unc5BiECko* retinas from arteries and capillaries that converted to a Claudin-5-/PLVAP+ phenotype (**Fig. 2o**).

### Endothelial Unc5B regulates Norrin/β-catenin signaling

As Claudin-5 and PLVAP are known targets of Norrin/β-catenin signaling at the BRB, we tested if endothelial Unc5B regulated Norrin/β-catenin signaling. Compared to TAM-treated cre-negative littermate controls, western blot of protein lysates from *Unc5BiECko* retinas showed reduced phosphorylation levels of LRP5, which is essential for proper Norrin/β-catenin signaling at the BRB (**Fig. 3a-c**), while phosphorylation levels of the Wnt/β*-*catenin co-receptor LRP6 were similar in retinal lysates from controls and mutants (**Supp. Fig. 3a-c**). Expression levels of β-catenin and of the transcriptional effector LEF1 were also reduced in *Unc5BiECko* retina lysates compared to controls (**Fig. 3a-c, Supp. Fig. 3d**.).

**Figure 3:**
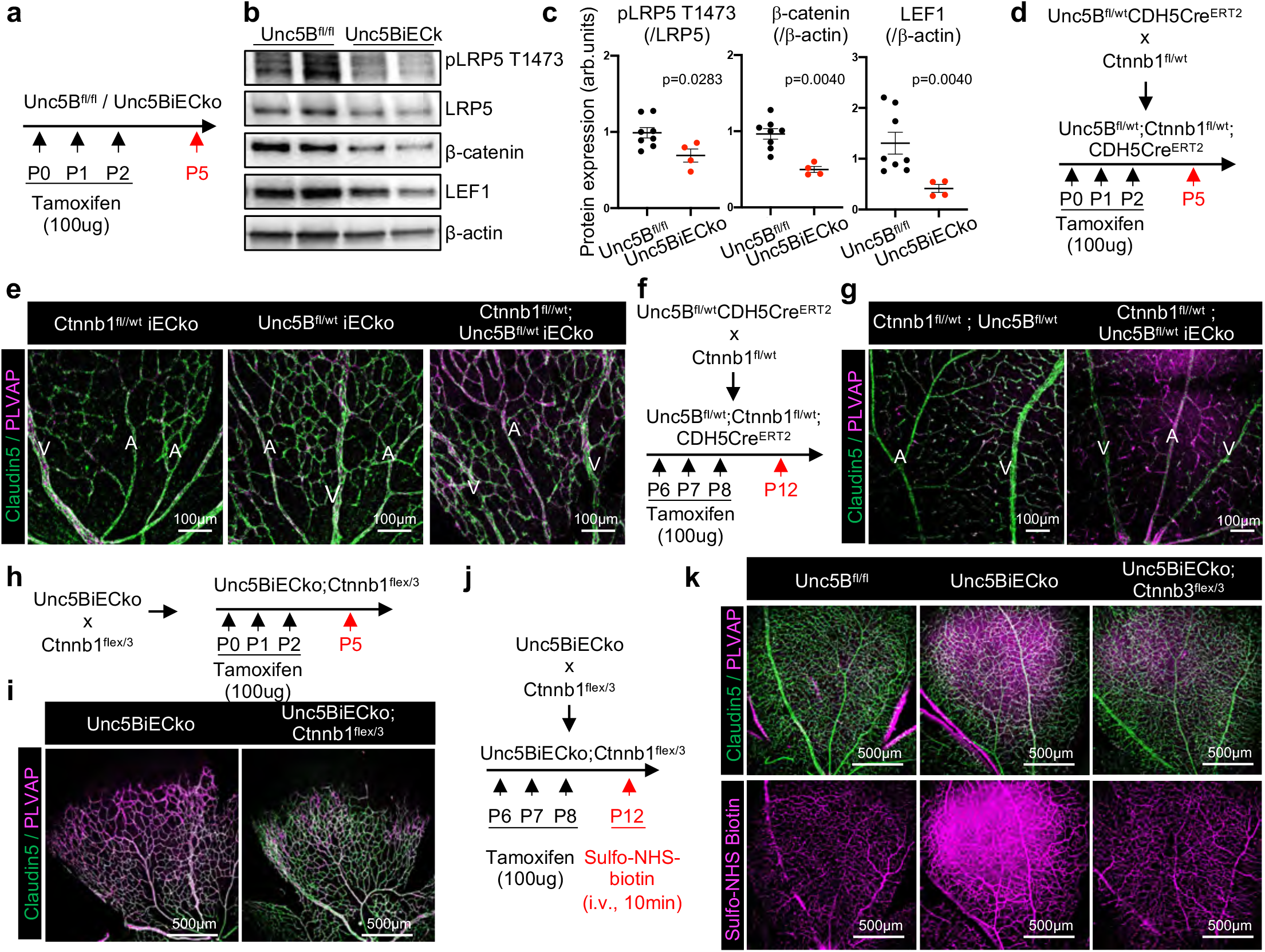
Endothelial Unc5B regulates BRB permeability via Norrin/β-catenin signaling. (a) *Unc5B* gene deletion strategy using tamoxifen injection in P5 neonates. (b,c) Western blot (b) and quantification (c) of P5 retina protein extracts. (d) *Unc5B* and *Ctnnb1* gene deletion strategy using tamoxifen injection in P5 retinas. (e) Immunofluorescence staining and confocal imaging on whole-mount P5 retinas with the indicated antibodies. (f) *Unc5B* and *Ctnnb1* gene deletion strategy using tamoxifen injection in P12 retinas. (g) Immunofluorescence staining and confocal imaging on whole-mount P12 retinas with the indicated antibodies. (h) *Unc5B* gene deletion and *Ctnnb1*^*flex/3*^ gene overexpression strategy using tamoxifen injection in P5 retinas. (i) Immunofluorescence staining and confocal imaging on whole-mount P5 retinas with the indicated antibodies. (j) *Unc5B* gene deletion and *Ctnnb1*^*flex/3*^ gene overexpression strategy using tamoxifen injection in P12 retinas. (k) Immunofluorescence staining and confocal imaging on whole-mount P12 retinas with the indicated antibodies, 10min after i.v. sulfo-NHS-biotin injection. All data are shown as mean+/-SEM. Two-sided Mann-Whitney U test was performed for statistical analysis.

To test for genetic interaction between *Unc5B* and *β-catenin*, we generated TAM-inducible endothelial-specific heterozygous loss of *Unc5B* (*Unc5B*^*fl/WT*^*iECko* mice) or *β-catenin* (*Ctnnb1*^*fl/WT*^*iECko* mice). *Unc5B*^*fl/WT*^*iECko and Ctnnb1*^*fl/WT*^*iECko* single heterozygote P5 and P12 retinas showed a BRB-competent Claudin-5+/PLVAP-expression pattern (**Fig. 3d-g**), revealing that a single allele of *Unc5B* or *β-catenin* was sufficient to maintain BRB integrity. However, double heterozygous *Unc5B*^*fl/WT*^;*Ctnnb1*^*fl/WT*^*iECko* P5 and P12 retinas exhibited conversion of arteries and capillaries to a Claudin-5-/PLVAP+ state (**Fig. 3d-g**), demonstrating that *Unc5B* and *β-catenin* genetically interact to maintain BRB ECs in a Claudin-5+/PLVAP-state.

Next, we overexpressed an activated form of β-catenin by crossing *Unc5BiECko* with *Ctnnb1*^*flex/3*^ mice^57^ (**Fig. 3h**), which enhances Norrin/β-catenin signaling *in vivo*^*13,58*^. The resulting offspring (*Unc5BiECko*; *Ctnnb1*^*flex/3*^) displayed decreased PLVAP protein expression, along with increased Claudin-5 protein expression compared to *Unc5BiECko* retinas at P5 (**Fig. 3h,i**) and P12 (**Fig. 3j,k**). Moreover, i.v. injection of sulfo-NHS-biotin into *Unc5BiECko; Ctnnb1*^*flex/3*^ mice showed reduced BRB leakage compared to *Unc5BiECko* mice (**Fig. 3j.k**).

We investigated other signaling pathways that might be implicated in Unc5B-mediated BRB leakage such as SOX17, a transcription factor that mediates arterial induction and maintenance of BBB integrity^59,60^. SOX17 was specifically expressed by arterial ECs in the retina of control littermates and *Unc5BiECko* retinas (**Supp. Fig. 3d**). Next, we examined Dll4 which regulates arterial BRB integrity by suppressing transcytosis and Caveolin1^61^ Compared to TAM-treated cre-negative littermate control, *Unc5BiECko* retinas did not show significant changes in DLL4 or Caveolin1 expression levels (**Supp. Fig. 3e**). These results suggest that Unc5B regulates BRB integrity mainly via Norrin/β-catenin signaling and independently of SOX17, DLL4 or Caveolin1.

### Netrin-1 regulates BRB development and integrity

To determine whether Unc5B ligands Netrin-1 and Robo4 regulated BRB integrity and Norrin/β-catenin pathway activation in retina ECs, we generated inducible *Netrin-1* global KO mice (hereafter *Ntn1iko*) by crossing *Netrin-1*^*fl/fl*^ mice with a *ROSA26Cre*^*ERT2*^ driver (Jackson laboratories) to induce ubiquitous gene deletion upon TAM injection. Compared to TAM-treated cre-negative littermate controls, P5 *Ntn1iko* retinas displayed increased vascular density and no change in outgrowth (**Fig 4a-c**). *Ntn1iko* retinas revealed conversion of arteries and capillaries to a leaky Claudin-5-/PLVAP+ phenotype similar to *Unc5BiECkos* (**Fig. 4d,e**), while *Robo4* global KO mice did not exhibit any EC conversion and remained in a BRB-competent Claudin-5+/PLVAP-state (**Supp. Fig. 4a,b**). We previously generated a monoclonal anti-Unc5B-3 antibody that binds human, rat and mouse Unc5B with high affinity and specifically blocks Netrin-1 binding to Unc5B^*51*^. I.p. injection of anti-Unc5B-3 (10mg/kg) for 2h followed by i.v. injection of sulfo-NHS-biotin revealed BRB breakdown compared to CTRL IgG injected littermate controls (**Fig. 4f,g**), demonstrating that blocking Netrin-1 binding to Unc5B is sufficient to open the BRB on-demand. Western blot from *Ntn1iko* retina protein lysates confirmed *Netrin-1* deletion, along with decreased expression of pLRP5, β-catenin, Claudin-5 and increased PLVAP expression (**Fig. 4h-j**), suggesting that Netrin-1 binding to Unc5B regulates Norrin/β-catenin signaling at the BRB. As *Ntn1* mRNA expression was virtually absent from the retina (**Supp. Fig. 1c**), we speculate that the relevant ligand source may come from the circulation.

**Figure 4:**
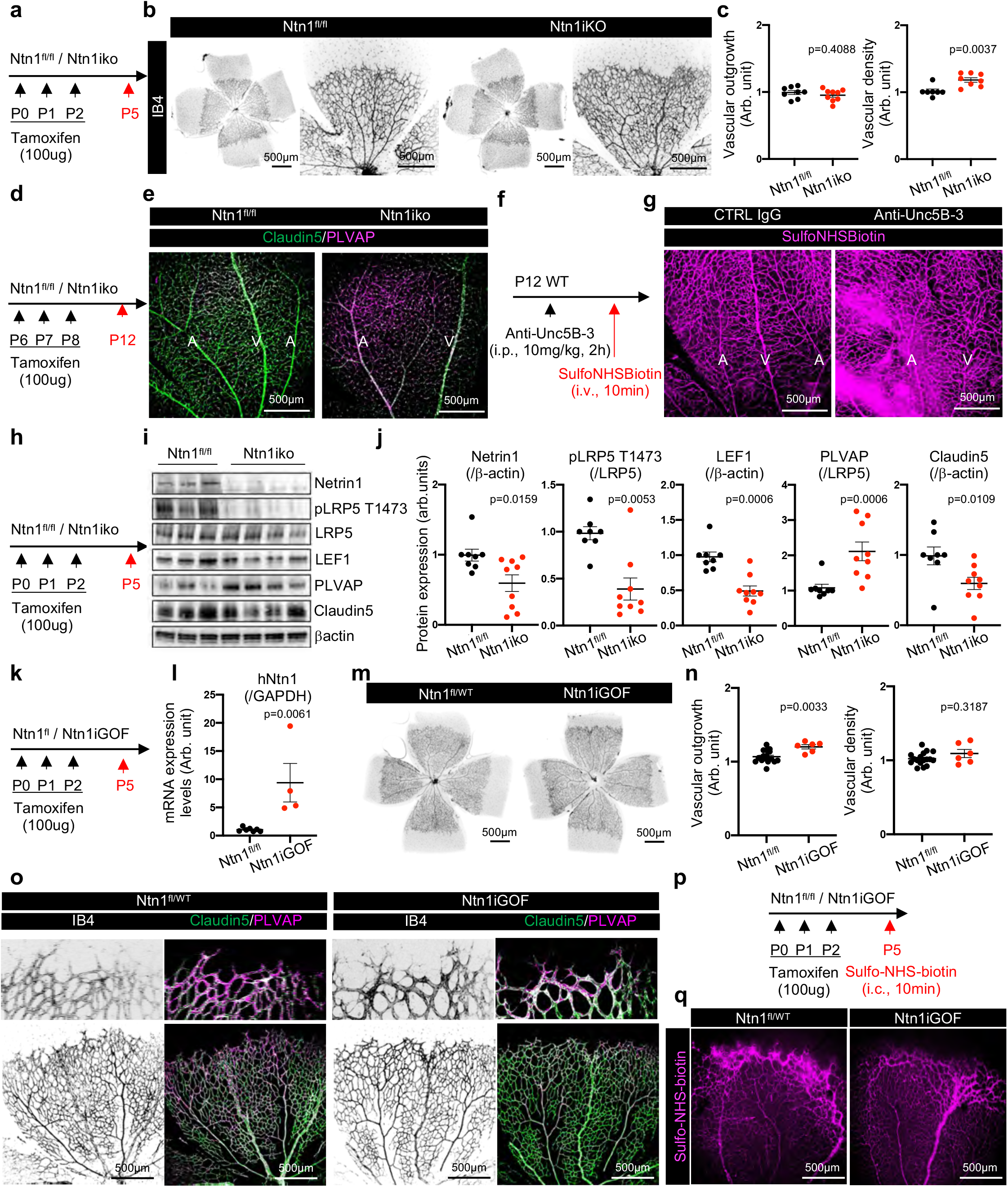
Netrin-1 regulates BRB development and integrity. (a) *Ntn1* gene deletion strategy using tamoxifen injection in P5 neonates. (b) IB4 immunofluorescence staining and confocal imaging on whole-mount P5 retinas. (c) Quantification of IB4+ vascular outgrowth and density in P5 retinas. (d) *Ntn1* gene deletion strategy using tamoxifen injection in P12 neonates. (e) Immunofluorescence staining and confocal imaging on whole-mount P12 retinas with the indicated antibodies. (f) Netrin1 binding to Unc5B blockade strategy using anti-Unc5B-3 antibody. (g) Confocal imaging on P12 retinas i.p. injected with anti-Unc5B-3 (2h, 10mg/kg) followed by i.v. sulfo-NHS-biotin injection for 10min. (h) *Ntn1* gene deletion strategy using tamoxifen injection in P5 neonates. (i,j) Western blot (i) and quantification (j) of P5 retina protein extracts. (k) *Ntn1* gene overexpression strategy using tamoxifen injection in P5 neonates. (l) qPCR analysis of P5 retina mRNA extracts. (m) IB4 immunofluorescence staining and confocal imaging on whole-mount P5 retinas. (n) Quantification of IB4+ vascular outgrowth and density in P5 retinas. (o) Immunofluorescence staining and confocal imaging on whole-mount P5 retinas with the indicated antibodies. (p) *Ntn1* gene overexpression strategy using tamoxifen injection in P5 neonates. (q) Confocal imaging on P5 retinas i.v. injected with sulfo-NHS-biotin for 10mim. All data are shown as mean+/-SEM. Two-sided Mann-Whitney U test was performed for statistical analysis.

To test if Netrin-1 overexpression would increase BRB properties *in vivo*, we generated TAM-inducible Netrin-1 gain of function mice (hereafter *Ntn1iGOF*) by intercrossing *loxStoplox;hNtn1*^62^ with *ROSACre*^*ERT2*^ driver to induce ubiquitous overexpression of human Netrin-1, which crossreacts with mouse receptors^43,62^. Gene overexpression was induced by TAM injections into neonates between P0-2 (**Fig 4k**), and qPCR analysis of P5 *Ntn1iGOF* mouse retina lysates revealed efficient Netrin-1 overexpression compared to TAM-treated cre-negative littermate controls (**Fig 4l**). *Ntn1iGOF* retinas showed no difference in vascular density compared to littermate controls, but displayed a slight but significant increase in vascular outgrowth (**Fig 4m,n**). To determine if *Ntn1GOF* increased BRB properties, we focused on P5 retina vascular front composed of Claudin-5-/PLVAP+ leaky superficial tip cells S compared to P12 BRB-competent diving tip cells (**Supp. Fig. 4c,d**). Compared to TAM-treated cre-negative littermate controls, *Ntn1GOF* P5 retinas exhibited conversion of ECs of the vascular front from a Claudin-5-/PLVAP+ leaky state to a Claudin-5+/PLVAP-phenotype (**Fig. 4o**,). Additionally, i.v. injection of sulfo-NHS-biotin into *Ntn1iGOF* mice showed reduced BRB leakage at the angiogenic front in P5 retinas, compared to TAM-injected cre-negative littermate controls (**Fig. 4p,q**).

### Mural cell Unc5B contributes to retinal angiogenesis and BRB regulation

To study Unc5B function in mural cells, and to address possible Unc5B functions in other retinal cells, we generated two additional mouse lines carrying pericyte-specific *Unc5B* deletion and global *Unc5B* deletions. Pericyte *Unc5B* knockout mice (hereafter *Unc5BiPCko)* were generated by intercrossing *Unc5B*^*flfl* 51^ and *PDGFRβCre*^*ERT2* 63^ mice, and global *Unc5B* knockout mice (hereafter *Unc5Biko)* were generated by intercrossing *Unc5B*^*flfl*^ and *RosaCre*^*ERT2*^ (Jackson Laboratories).

Gene deletion was induced by TAM injections into neonates between P0-2, and immunostaining and western blot analysis of retinal lysates revealed efficient Unc5B deletion in *Unc5Biko* retinas (**Supp. Fig 5a-c**). In the *Unc5BiPCko* mice, immunostaining with an anti-Unc5B antibody revealed reduced protein expression in pericytes while EC Unc5B expression was maintained (**Supp. Fig. 5d**), confirming mural cell-specific deletion. By Western blot, Unc5BiPCko retinal lysates maintained Unc5B expression at levels slightly higher when compared to controls, although this did not reach statistical significance (**Supp. Fig. 5e,f**). However, qPCR on *Unc5BiPCko* isolated purified brain pericytes demonstrated efficient Unc5B gene deletion compared to TAM-treated cre-negative littermate controls (**Supp. Fig. 5g,h**).

**Figure 5:**
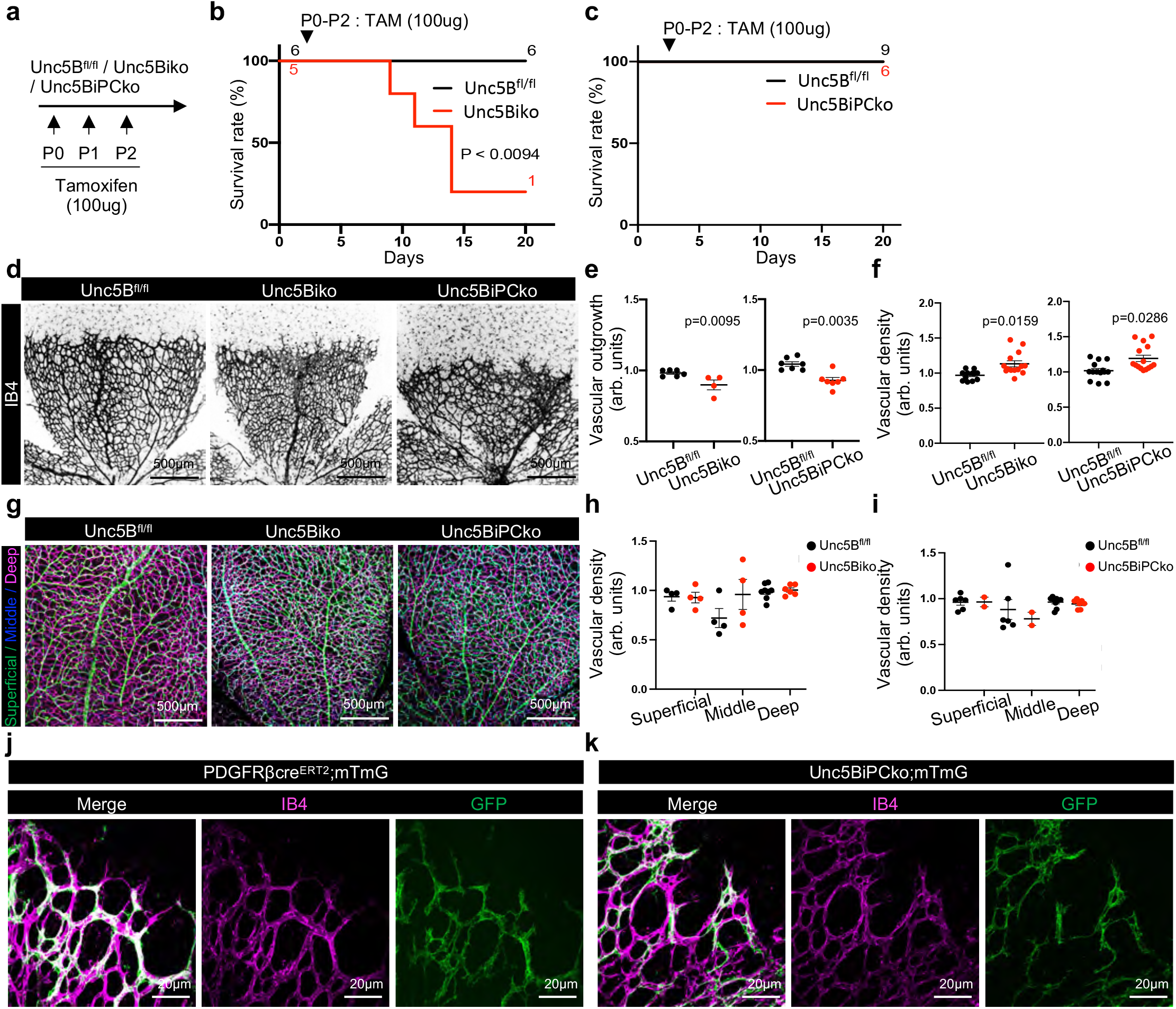
Mural cell Unc5B contributes to retinal angiogenesis. (a) Global or Pericyte *Unc5B* gene deletion strategy using tamoxifen injection. (b,c) Survival curve after neonatal global (b) or pericyte (c) *Unc5B* gene deletion. (d) Immunofluorescence staining and confocal imaging on whole-mount P5 retinas with the indicated antibodies. (e,f) Quantification on IB4+ vascular outgrowth and density in P5 retinas. (g) Immunofluorescence staining of IB4+ vessel and confocal imaging on whole-mount P12 retinas. Superficial, intermediate and deep plexus were color-coded. (h,i) Quantification of superficial, intermediate and deep plexus vascular density in P12 retinas. (j,k) Immunofluorescence staining and confocal imaging on the vascular front of whole-mount P5 retinas with the indicated antibodies. All data are shown as mean+/- SEM. Mantel-cox test was performed for survival curve statistical analysis. Mann-Whitney U test was performed for statistical analysis between two groups. ANOVA followed by Bonferroni’s multiple comparisons test was performed for statistical analysis between multiple groups.

Neonatal TAM injections induced lethality of *Unc5Biko* mice (**Fig. 5a,b**) as with *Unc5BiECko* mice^51^, but *Unc5BiPCko* mice survived and appeared healthy (**Fig. 5c**). When gene deletion was induced at P0-2 and retinal vasculature examined at P5, we found that both, *Unc5Biko* and *Unc5BiPCko* mice displayed transient defects in angiogenesis, with decreased vascular outgrowth and increased vascular density at P5 (**Fig 5d-f**) that were no longer observed when gene deletion was induced at P6-8 and retinas examined at P12 (**Fig. 5g-i)**. Hence, both endothelial and pericyte Unc5B transiently modulate vascular development of the superficial plexus. Interestingly, analysis of P5 *Unc5BiPCko* mice crossed with mTmG reporter mice^64^ revealed pericyte covered superficial tip cells in the *Unc5BiPCko* mice (**Fig. 5j,k**) as a potential cause for their dysregulated angiogenesis. This demonstrates that both endothelial and pericyte Unc5B regulate angiogenesis of the superficial vascular plexus.

To assess BRB permeability, we examined leakage of intracardiac injected sulfo-NHS-biotin in P12 mice after gene deletion between P6 and P8 (**Fig. 6a**). *Unc5Biko* mice exhibited sulfo-NHS-biotin leak, although to a lesser extent than *Unc5BiECko* mice (**Fig. 6b)**. By contrast, the tracer remained confined to the vessels in CTRL and *Unc5BiPCko* mice (**Fig. 6b**). Tracer leak experiments in *Unc5BiPCko* and *Unc5Biko* mice analyzed at P5 demonstrated similar results (**Supplementary Fig. 6a-d**). Some ECs in the *Unc5Biko* mice converted to a PLVAP+, Claudin5 low phenotype, but as with tracer leak, the effect was less pronounced mice when compared to the *Unc5BiECko* mice (**Fig. 6c,d**). Interestingly, *Unc5BiPCko* mice showed robust Claudin-5 immunofluorescence when compared to controls (**Fig. 6c,d**), and western blot analysis confirmed increased Claudin5 levels in *Unc5BiPCko* retinal lysates, although this did not reach statistical significance (**Fig. 6e-g**). *Unc5Biko* and *Unc5BiPCko* showed no or minimal change in astrocyte and pericyte coverage, determined by GFAP and PDGFRβ immunostaining and western blot analysis, respectively (**Supplementary Fig. 6e-j**). *Unc5BiPCko* retinal lysates also had slightly increased levels of pLRP5 T1473, but this effect did not translate to alterations in β-catenin or LEF1 expression (**Fig. 6e-g**). These data suggest that pericyte Unc5B somehow opposes endothelial Unc5B signaling, and that global Unc5B deletion showed effects of both, endothelial and pericyte Unc5B deletion. qPCR analysis on endothelial cells extracted from *Unc5BiPCko* brains revealed increased levels of *UNC5B* and *CLDN5* compared to littermate controls (**Supp. Fig. 6k,l**), suggesting a possible mechanism whereby PC Unc5B signals to limit endothelial Unc5B expression and BRB signaling.

**Figure 6:**
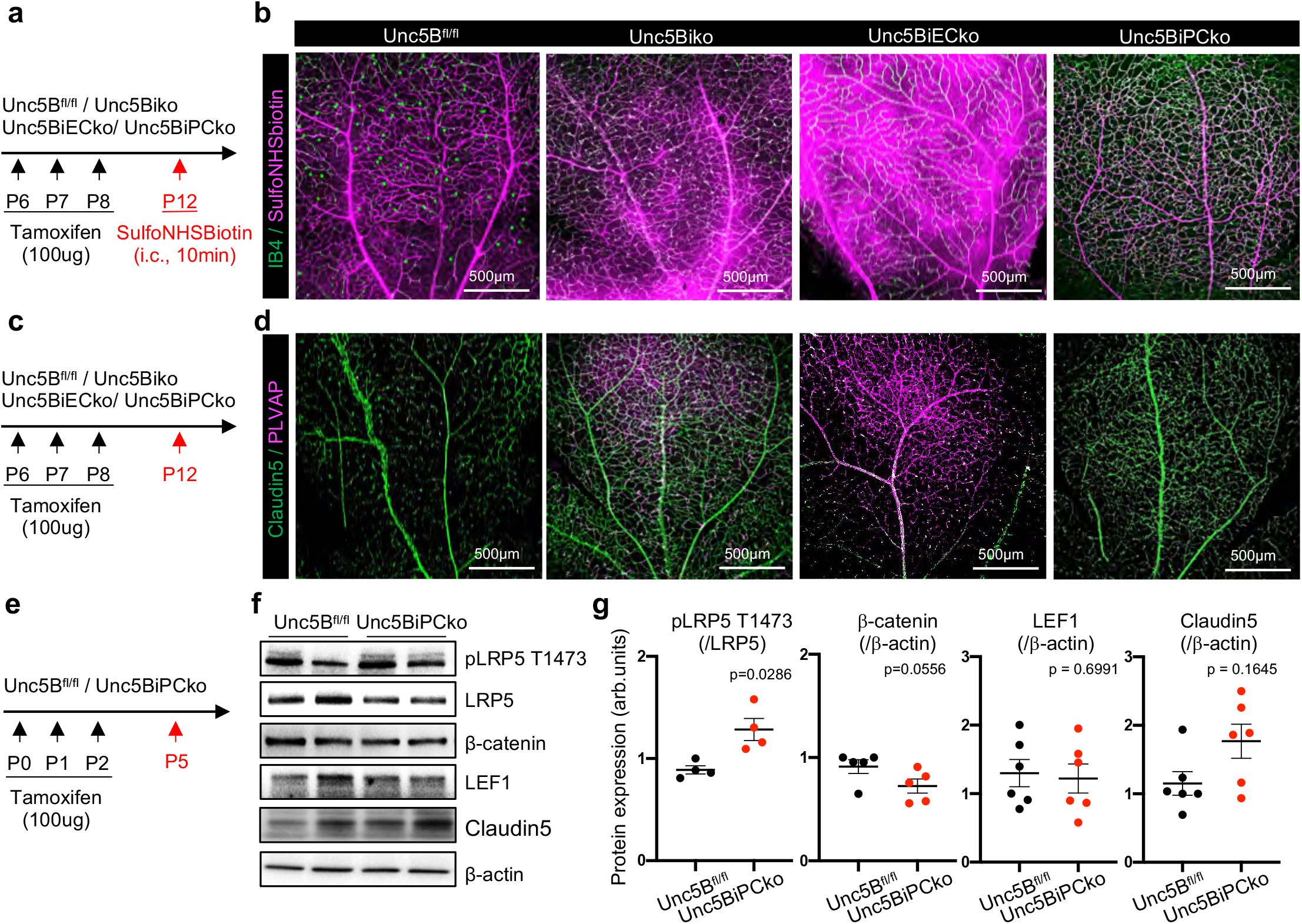
Mural cell Unc5B contributes to BRB regulation. (a) *Unc5B* gene deletion strategy using tamoxifen injection in P12 neonates. (b) Immunofluorescence staining and confocal imaging on whole-mount P12 retinas with the indicated antibodies, 10min after i.v. sulfo-NHS-biotin injection. (c) *Unc5B* gene deletion strategy using tamoxifen injection in P12 neonates. (d) Immunofluorescence staining and confocal imaging on whole-mount P12 retinas with the indicated antibodies. (e) *Unc5B* gene deletion strategy using tamoxifen injection in P5 neonates. (f,g) Western blot (f) and quantification (g) of P5 retina protein extracts. All data are shown as mean+/-SEM. Mann-Whitney U test was performed for statistical analysis between two groups.

### Netrin-1 and Norrin cooperate to induce β-catenin signaling

To determine whether Netrin-1 binding to Unc5B cooperated with Norrin at the BRB, we treated serum-starved primary mouse brain ECs with Netrin-1, Norrin, or with a combination of both Netrin-1 and Norrin at different concentrations (**Fig. 7a-c**). Stimulation with either Netrin-1 or Norrin induced LRP5 phosphorylation with a peak between 5 to 30min (**Fig. 7a-c**). However, combined stimulation with Netrin-1 and Norrin led to increased and sustained LRP5 phosphorylation levels when compared to either ligand alone, with a peak at 1h **(Fig. 7a-c)**. Netrin-1 stimulation also induced transient phosphorylation of the BBB Wnt/β-catenin co-receptor LRP6 as previously described^51^ (**Supp. Fig. 7a,b**). Norrin stimulation induced lower levels of pLRP6 when compared to pLRP5, and combination of both Netrin-1 and Norrin did not show additive effects on LRP6 activation **(Supp. Fig. 7a,b**). To determine whether increased phosphorylation of LRP5 led to increased β-catenin-mediated transcription, primary mouse brain ECs were transfected with a luciferase construct containing four native TCF/LEF binding sites (TOPflash) or its negative-control counterpart (FOPflash) containing four mutated LEF/TCF binding sites (**Fig. 7d**). Mouse brain ECs treated with Netrin-1 or Norrin alone revealed increased luciferase activity starting 1h after stimulation and with a maximal response after 8h, demonstrating that Netrin-1 and Norrin-mediated LRP5 activation increased β-catenin transcriptional activity (**Fig. 7d**). Combinatory stimulation of Netrin-1 and Norrin induced a more potent luciferase activity compared to Netrin1 or Norrin alone, revealing additive activation of β-catenin-mediated transcription (**Fig. 7d**).

**Figure 7:**
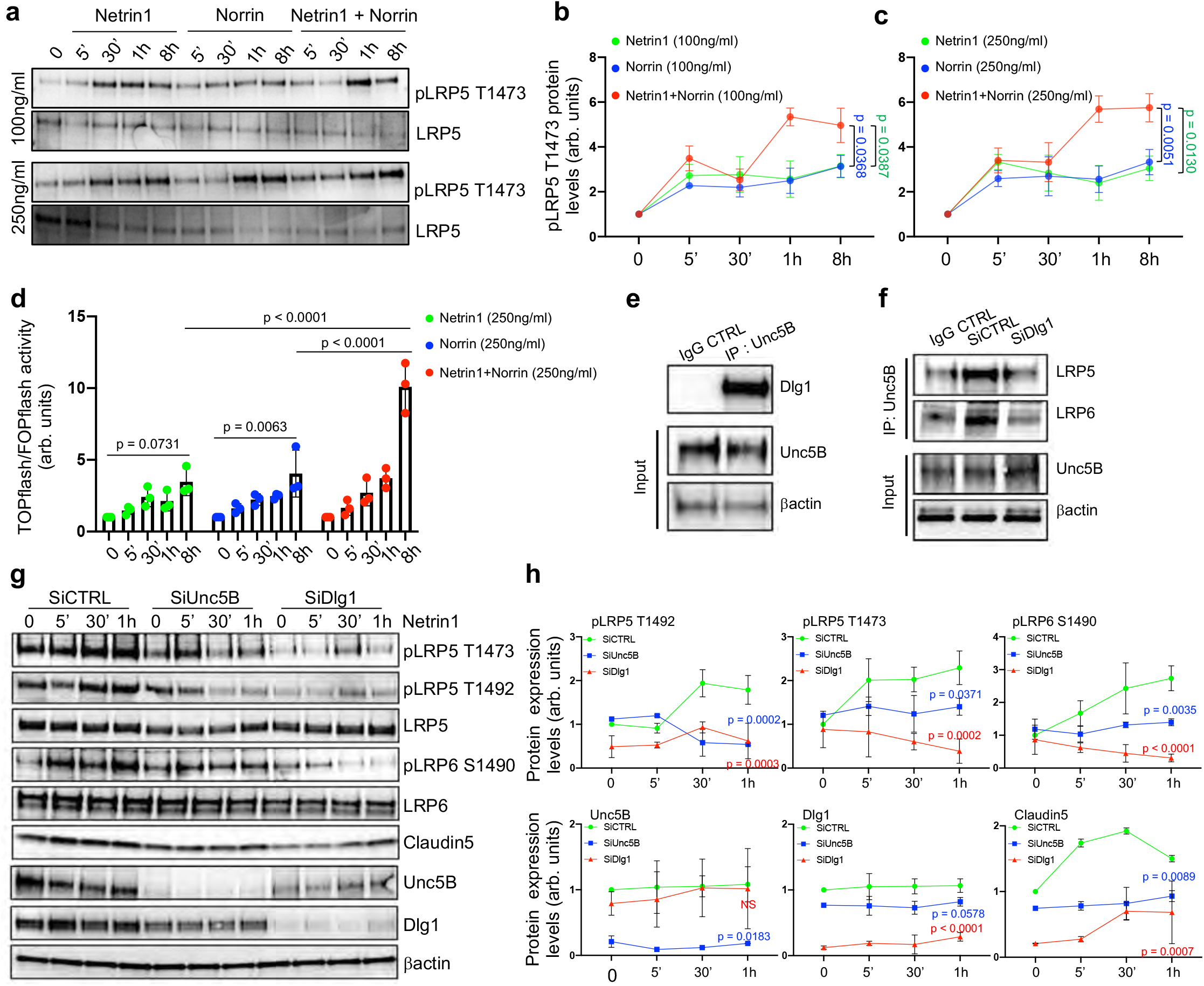
Netrin-1 and Norrin cooperate to induce β-catenin signaling (a-c) Western blot (a) of mouse brain EC protein extracts treated for the indicated times (min) with Netrin-1, Norrin, or a combination of both. (b,c) Quantification of LRP5 phosphorylation levels over time (d) Mouse brain ECs were transfected with the β-catenin transcriptional luciferase reporter TOPflash or FOP flash control and stimulated with Netrin-1, Norrin, or a combination. Cells were harvested in 1x cell culture lysis reagent and luciferase activity was quantified using a plate-based luminometer. (e) CTRL IgG and Unc5B immunoprecipitation from cultured mouse brain ECs. (f) CTRL IgG and Unc5B immunoprecipitation on CTRL or Dlg1 SiRNA-treated mouse brain ECs. (g,h) Western blot (g) and quantification (h) of CTRL, Unc5B or Dlg1 SiRNA-transfected mouse brain ECs treated with Netrin-1 (250ng/ml). All data are shown as mean+/-SEM. ANOVA followed by Bonferroni’s multiple comparisons test was performed for statistical analysis between multiple groups.

Mechanistically, we tested if the scaffolding protein Dlg1 interacted with Unc5B and mediated Netrin-1 activation of LRP5-T1473, LRP5-T1492 and LRP6-S1490 phosphorylation. Unc5B immunoprecipitation of primary mouse brain ECs pulled down Dlg1 (**Fig. 7e**). *Dlg1* siRNA-mediated knockdown reduced pulldown of LRP5 and LRP6 with Unc5B compared to Ctrl siRNA treated cells (**Fig. 7f**), demonstrating that Dlg1 association with Unc5B regulated complex formation of Unc5B with LRP5. Moreover, Netrin-1-treated serum-starved CTRL, *Unc5B* or *Dlg1* SiRNA transfected ECs cells revealed that both Unc5B and Dlg1 are essential for Netrin-1-induced LRP5 and LRP6 phosphorylation (**Fig. 7g,h**). Netrin-1 stimulation on CTRL cells also induced expression of Claudin-5, attesting β-catenin-mediated transcription activity, which was reduced upon *Unc5B* and *Dlg1* SiRNA treatment (**Fig. 7g,h**).

## DISCUSSION

Our data reveal Netrin-1 signaling via endothelial Unc5B as a novel regulator of BRB development. We observed that TAM-inducible endothelial-specific deletion of *Unc5B*, or global inducible *Ntn1* deletion led to BRB leakage mainly from retinal arteries and capillaries, which converted from a Claudin-5+/PLVAP-state to a Claudin-5-/PLVAP+ phenotype. As Unc5B is predominantly expressed in retinal arteries and capillaries, these data support that Netrin-1 signals through endothelial Unc5B to induce a BRB-competent state in these cells. Conversely, genetic Ntn1 overexpression converts leaky vessels at the P5 angiogenic front to a non-leaky Claudin-5+/PLVAP-state. As negligible Netrin-1 was present in retinal scRNAseq data, this Netrin-1 may come from the circulation, produced by the heart, lungs, trachea or adipose tissue^65^. Taken together with our previous demonstration of Netrin-1-Unc5B BBB permeability regulation^51^ the data reveal endothelial Unc5B signaling as a CNS-wide endothelial barrier permeability control point that can be manipulated to decrease or increase barrier properties on demand.

Mechanistically, retinal Unc5B maintains LRP5 phosphorylation at T1473 and T1492. These phosphorylation sites provide a docking site for the adapter protein Axin1, which inhibits the β-catenin destruction complex and promotes nuclear β-catenin translocation, activation of Norrin-induced-LEF1 signaling and BRB gene expression regulation^17,66^. We show genetic interaction between *Unc5B* and *Ctnnb1* in retinal ECs, in that heterozygous deletions in either *Unc5B* or *β-catenin* displayed BRB competent Claudin-5-+/PLVAP-ECs while combined heterozygous deletions led to conversion to a leaky Claudin-5-/PLVAP+ phenotype at the BRB. Conversely, endothelial-specific β-catenin overexpression in *Unc5BiECko* mice rescued both ECs conversion to a BBB-competent state and leakage to i.v. injected tracer, demonstrating that Unc5B regulates BRB integrity through its regulation of Norrin/β-catenin signaling. *In vitro*, we observed that combined stimulation of ECs with Netrin-1 and Norrin additively regulated LRP5 phosphorylation and TOPflash gene expression. These data suggest that Unc5B cooperates with Norrin to maintain β-catenin activity above the specific threshold required to maintain BRB ECs in a Claudin-5+/PLVAP-state^13,17,19,20,22^. Because β-catenin signaling was decreased in *Ntn1* and *Unc5B* mutants, while Unc5B expression was unaltered in loss or gain of endothelial *Ctnnb1* mutants^51^, the data is consistent with Netrin-1-Unc5B signaling acting as an upstream regulator of the BBB and BRB.

Unc5B interaction with LRP5 required the scaffolding protein Dlg1, which was previously shown to maintain the BRB and the BBB^30^. Loss of endothelial Dlg1 function converted retina and brain ECs to a leaky, Claudin-5-/PLVAP+ state, and *Dlg1* genetically interacted with *Ndp, Fz4* and *Tspan* to regulate BRB and BBB integrity, via interaction with proteins other than Frizzled receptors^30^. We could co-immunoprecipitate Unc5B with Dlg1, and we show that siRNA-mediated *Dlg1* knockdown in brain ECs decreased Unc5B pulldown of LRP5 and activation of LRP5 and LRP6 phosphorylation by Netrin-1. Hence, our data suggest that Dlg1 enhances both Wnt- and Norrin-β-catenin signaling via Unc5B, providing mechanistic insight into pathway convergence. Further experiments will need to clarify how this interaction occurs, but we speculate that Dlg1 binds to the intracellular Unc5B UPA domain, which is required for its interaction with LRP6^51^.

Despite the evidence that BRB gene regulation of Unc5B depended on β-catenin, we observed differences between both pathways. Endothelial-specific *Ctnnb1* deletion induced Claudin-5-/PLVAP+ conversion in veins and capillaries^13^, while loss of *Unc5B* converted arteries and capillaries, but not veins. *Lrp5*^*–/–*^ *Lrp6*^*CKO/+*^ *Tie2Cre* retinas converted all ECs to a leaky Claudin-5-/PLVAP+ phenotype^13^. The observed differences in arteriovenous zonation deserve further investigation and could be due to the involvement of β-catenin-independent Unc5B signaling pathways in arteries. However, our data shows that Unc5B regulates arterial BRB integrity in the absence of changes in expression of signaling pathways that regulate arterial BRB integrity such as Sox17, Dll4 and Caveolin1^59-61^.

Another interesting difference concerned retinal angiogenesis regulation by Unc5B and canonical Norrin-β-catenin signaling. *Unc5B* deletion did not affect formation of the deeper vascular layers, while *Ndp, Fz4, Lrp5* and *Ctnnb1* mutation prevents formation of these vessels^13,22,54,67^. We speculate that *Unc5B* deletion maintains β-catenin levels sufficiently high to induce vessel diving, but not high enough to maintain BRB gene expression. In support of this idea, we show that global *Unc5b* mutants display less BRB leak when compared to *Unc5BiECKo*, while pericyte *Unc5B* deletion tended to enhance EC Unc5B expression levels, thereby increasing retinal LRP5 activation and tending towards enhanced BRB function. The opposing actions between EC and PC Unc5B may explain the lesser extent of BRB leak observed in global *Unc5B* mutants, and the existence of this regulation supports that endothelial Unc5B signaling levels are carefully controlled during BRB development. In line with this idea, dysregulation of Netrin-1 expression in human patients with diabetic retinopathy was shown to contribute to vascular leak and macular edema^68^.

Unc5B and Norrin signaling also both regulate angiogenesis of the superficial vascular layer and S-tip cell sprouting. The angiogenic front of the superficial plexus remains leaky despite expression of Unc5B and β-catenin, probably due to additional permeability regulators such as VEGF which is expressed at high levels in astrocytes ahead of the vascular front. The sprouting angiogenesis defects in Unc5B deficient P5 retinas were not rescued by *Ctnnb1* overexpression, and likely depend on other pathways that remain to be identified. Our data show that regulation of superficial sprouting by Netrin-1-Unc5B is unexpectedly complex. *Unc5B* mutants display decreased vascular outgrowth and increased vascular density, consistent with previous observations that Netrin-1 repels tip cells via Unc5B^34,48^. Interestingly however, we found that mural cell Unc5B signaling contributed to sprouting defects, leading to increased pericyte coverage of angiogenic sprouts in *Unc5BiPCko* mice that may contribute to the observed sprouting defects. Moreover, while *Ntn1* mutants also displayed transient angiogenesis defects, these did not mirror the *Unc5BiECko* mutants. Because of the transient nature of the angiogenic defects, its underlying mechanisms were not further investigated at this point. Instead, we believe the permanent barrier damages induced by Ntn1-Unc5B loss of function demonstrate the importance of this pathway for vascular maturation and neuroprotection.

Recent studies showed that engineering of Wnt/Ndp signaling ligands into BBB-specific Wnt activators enable BBB repair in during brain tumors and ischemic stroke^69^. Other studies have developed agonists that elicit a β-catenin signaling response^70^ and that induce BRB stabilization and prevention of diabetic macular edema, including an agonistic antibody that activates the FZD4:NDP receptor^71^. Approaches to enhance Netrin-1 binding to endothelial Unc5B could synergize with such therapies and offer more broad application through the regulation of both the BBB and the BRB.

## MATERIALS AND METHODS

### Mouse models

All protocols and experimental procedures were approved by the Institutional Animal Care and Use Committee (IACUC). *Unc5B*^*fl/fl*^ mice^51^ were bred with *Cdh5Cre*^*ERT2*^ mice^56^ or *PDGFRβCre*^*ERT2* 63^ or ROSA*Cre*^*ERT2*^ (Jackson laboratories), βcatenin GOF *Ctnnb1*^*flex/3*^ mice, *Ctnnb1*^*fl/fl*^ mice, *Robo4*^*-/-*^ mice and *Ntn1*^*fl/fl*^ mice were described previously^10,48,50,57,72-74^. Inducible Netrin1 gain of function mice were generated by intercrossing *loxStoplox;hNtn1*^62^ with *ROSACre*^*ERT2*^ (Jackson laboratories) driver to induce ubiquitous overexpression of Netrin-1. Gene deletion or overexpression were induced by injection of tamoxifen (Sigma T5648) diluted in corn oil (Sigma C8267). Postnatal gene deletion was induced by 3 injections of 100ug of tamoxifen at P0, P1 and P2 for analysis at P5, or by 3 injections of 100ug of tamoxifen at P6, P7 and P8 for analysis at P12

### Cell culture

C57BL/6 Mouse Primary Brain Microvascular Endothelial Cells were purchased from Cell Biologics (C57-6023). Cells were cultured in Dulbecco’s Modified Eagle’s Medium (DMEM) high glucose (Thermo Fisher Scientific, 11965092) supplemented with 10% fetal bovine serum (FBS) and 1% Penicillin Streptomycin at 37 °C and 5% CO2 and split when confluent using Trypsin-EDTA (0.05%) (Life Technologies, 25300054). When indicated, cells were treated with recombinant Mouse Netrin-1 protein (R&D, 1109-N1-025), or with recombinant mouse Norrin protein (R&D, 3497-NR-025).

### Bioinformatics analysis

The scRNA-seq data from P6 and P10 control retinas were extracted from our previous work that is publicly available on NCBI’s Gene Expression Omnibus (accession no. GSE175895)^55^. Data processing and clustering were performed as previously described^55^. Gene expression analysis was conducted using the Seurat R-package (version 3.2.2)^75^.

### Western-Blot

Retinas were dissected and frozen in liquid nitrogen. They were lysed in RIPA buffer (Research products, R26200-250.0) supplemented with protease and phosphatase inhibitor cocktails (Roche, 11836170001 and 4906845001) using a Tissue-Lyser (5 times 5min at 30 shakes/second). For western blot on cell culture, cells were washed with PBS and lysed in RIPA buffer with protease and phosphatase inhibitors cocktails. All protein lysates were centrifuged 15min at 13200RPM at 4°C and supernatants were isolated. Protein concentrations were quantified by BCA assay (Thermo Scientific, 23225) according to the manufacturer’s instructions. 30ug of protein were diluted in Laemmli buffer (Bio-Rad, 1610747) boiled at 95°C for 5min and loaded in 4-15% acrylamide gels (Bio-Rad, 5678084). After electrophoresis, proteins were transferred on a polyvinylidene difluoride (PVDF) membrane and incubated in TBS 0.1% Tween supplemented with 5% BSA for 1hour to block non-specific binding. The following antibodies were incubated overnight at 4°C: Unc5B (Cell Signaling, 13851S), Netrin-1 (R&D, AF1109), Claudin-5 (Invitrogen, 35-2500), PLVAP (BD biosciences, 550563), PDGFRβ (Cell Signaling, 3169S), GFAP (DAKO, Z0334), pLRP6 (Cell Signaling, 2568S), LRP6 (Cell Signaling, 3395S), pLRP5 T1473 (Thermo Fisher Scientific, PA5105096), pLRP5 T1492 (Abcam, ab203306), LRP5 (Cell Signaling, #5731S), LEF1 (Cell Signaling, 2230S), βcatenin (Cell Signaling, 8480S), Dlg1 (Abcam, ab134156) and βactin (Sigma, A1978). Then, membranes were washed 4 × 10min in TBS 0.1% Tween and incubated with one of the following peroxidase-conjugated secondary antibodies diluted in TBS 0.1% Tween supplemented with 5%BSA for 2h at room temperature: horse anti-mouse IgG(H+L) (Vector laboratories, PI-2000), goat anti-rabbit IgG(H+L) (Vector laboratories, PI-1000), goat anti-rat IgG(H+L) (Vector laboratories, PI-9400). After 4 × 10min wash, western blot bands were acquired using ECL western blotting system (Thermo Scientific, 32106) or west femto maximum sensitivity substrate (Thermo Scientific, 34095) on a Biorad Gel Doc EQ System with Universal Hood II imaging system equipped with Image Lab software.

### Immunoprecipitation

Pierce™ protein A/G magnetic beads (Thermo Fischer, 88802) were washed 5 times 10min with RIPA buffer. 300ug of protein lysate were diluted in 1ml of RIPA buffer containing protease and phosphatase inhibitors and were incubated with 30ul of A/G magnetic beads for 1h at 4°C under gentle rotation. Protein lysates were harvested using a magnetic separator (Invitrogen) and were incubated overnight at 4°C under gentle rotation with 10ug of Unc5B antibody (R&D, AF1006) or control IgG. The next day, 40ul of protein A/G magnetic beads were added to each protein lysate for 2h at 4°C under gentle rotation. Beads were then isolated using a magnetic separator and washed 5 times with RIPA buffer. After the last wash, supernatants were removed and beads were resuspended in 40ul of Laemmli buffer (Bio-Rad, 1610747), boiled at 95°C for 5min and loaded onto 4-15% gradient acrylamide gels. Western blotting was performed as described above.

### Retina whole-mount immunostaining

Retinas were collected and placed in 3.7% formaldehyde at room temperature for 20minutes. Retinas were washed 3 × 10minutes with PBS and retinas were dissected and incubated with specific antibodies in blocking buffer (1% fetal bovine serum, 3% BSA, 0.5% Triton X-100, 0.01% Na deoxycholate, 0.02% Sodium Azide in PBS at pH 7.4) overnight at 4degree. The next day, retinas were washed 3 × 10minutes and incubated with secondary antibody for 2h at room temperature, then washed 3 × 10minute and post-fixed with 0.1% PFA for 10minutes and mounted using fluorescent mounting medium (DAKO, USA).

For mouse injected with fluorescent tracers, retinas were dissected in 3.7% formaldehyde to ensure dye fixation within the tissue during retina dissection.

The following antibodies were used: Unc5B (Cell signaling, 13851S), Claudin-5-GFP (Invitrogen, 352588), PLVAP (BD biosciences, 550563), GFAP (Millipore, MAB360), PDGFRβ (Cell Signaling, 3169S), CD13 (BD Biosciences, 558744), LEF1 (Cell Signaling, 2230S), SOX17 (R&D, AF1942), DLL4 (R&D, AF1389), Caveolin1 (Cell Signaling, D46G3), ERG (Abcam, ab92513), DAPI (Thermo Fischer, 62248). IB4 as well as all corresponding secondary antibodies were purchased from Invitrogen as donkey anti-primary antibody coupled with either Alexa Fluor 488, 568 or 647.

### Small interfering RNA knockdown experiments

Cells were transiently transfected with siRNA (Dharmacon). ON-TARGETplus Mouse Unc5b siRNAs (SMARTpool, L-050737-01-0005) were used for Unc5B gene deletion. ON-TARGETplus Mouse Dlg1 SiRNAs (SMARTpool, L-042037-00-0005 were used for Unc5B gene deletion. Transfection was performed using lipofectamine RNAi max (Invitrogen, 13778-075) according to the manufacturer’s instruction with siRNA at a final concentration of 25pmol in OptiMem for 8h. After transfection, cells were washed with PBS and fresh complete media was added for 48h.

### Quantitative real-time PCR analysis

mRNA was isolated using Trizol reagent (Life Technologies, 15596018) according to the manufacturer’s instructions and quantified RNA concentrations using nanodrop 2000 (Thermo Scientific). 300ng of RNA were reverse transcribed into cDNA using iScript cDNA synthesis kit (Bio-rad, 170-8891). Real-time qPCR was then performed in duplicates using CFX-96 real time PCR device (Bio-rad). Mouse GAPDH (QT01658692) was used as housekeeping gene for all reactions.

### Top flash experiment

Mouse brain ECs were transfected with the β-catenin transcriptional luciferase reporter TOPflash or FOP flash control. Transfection was performed using lipofectamine 3000 according to the manufacturer’s instruction. 48h post transfection, cells were starved for 16h and stimulated with Netrin-1, Norrin, or a combination of both. Then, cells were harvested in 1x cell culture lysis reagent (Promega E1531) for 10min at room temperature and luciferase activity was quantified using a Luciferase Assay System (Promega E1500) and a plate-based luminometer.

### Intravenous and intraperitoneal injection of tracer and antibodies

SulfoNHSbiotin tracer were injected intravenously in P12 mice at a concentration of 0.5 mg/g and left to circulate for 10min. For P5 neonates, Lysine-fixable Cadaverine conjugated to Alexa Fluor-555 (Invitrogen) was injected intraperitoneally at a concentration of 250ug cadaverine/20g of pups and left to circulate for 2h. CTRL IgG, and anti-Unc5B-3 were injected intraperitoneally at a concentration of 10mg of antibodies/kg of mice for 2h

### Confocal microscopy and image analysis

Confocal images were acquired on a laser scanning fluorescence microscope (Leica SP8) using the LASX software. 10X, 20X and 63X oil immersion objectives were used for acquisition using selective laser excitation (405, 488, 547, or 647 nm). Quantification of vascular outgrowth as well as was quantification of pixel intensity were performed using the software Image J. Vascular density was quantified with the software Angiotool by quantifying the vascular surface area normalized to the total surface area. Quantification of pixel intensity was performed using the software ImageJ.

### Statistical analysis

All *in vivo* experiments were done on littermates with similar body weight per condition. Statistical analysis was performed using GraphPad Prism 8 software. Mann-Whitney U test was performed for statistical analysis of two groups. ANOVA followed by Bonferroni’s multiple comparisons test was performed for statistical analysis between 3 or more groups. Mantel-cox test was performed for survival curve statistical analysis.

## FIGURE LEGENDS

**Supplemental Figure 1:**
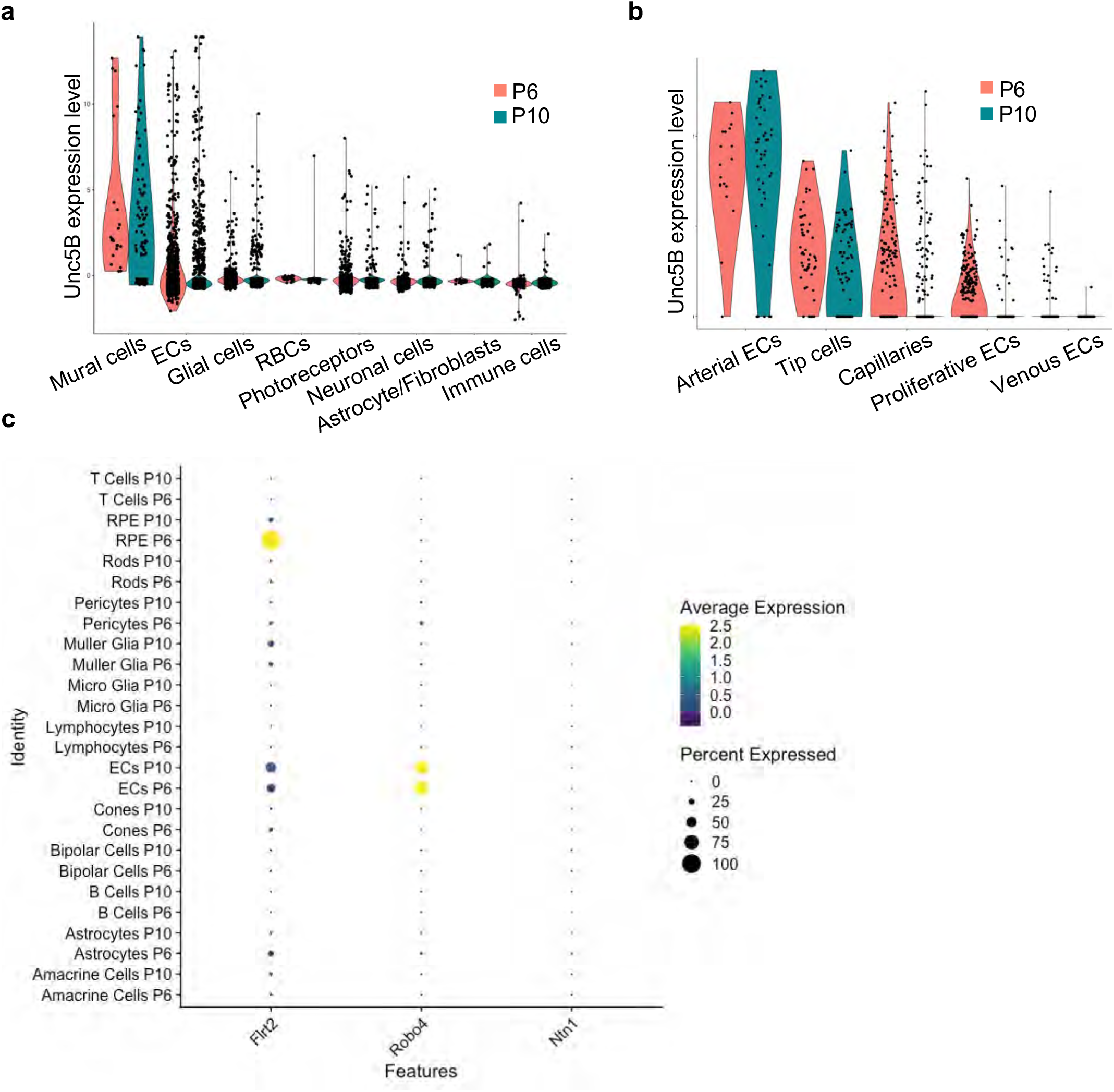
(a,b) Violin plots of the expression of Unc5B in retina cells^55^ (a) or in endothelial cell subclusters (b)^55^. (c) Dot plot expression levels and frequency of Unc5B ligands among retina cell clusters^55^. Color scale: yellow: high expression; dark blue: low expression.

**Supplemental Figure 2:**
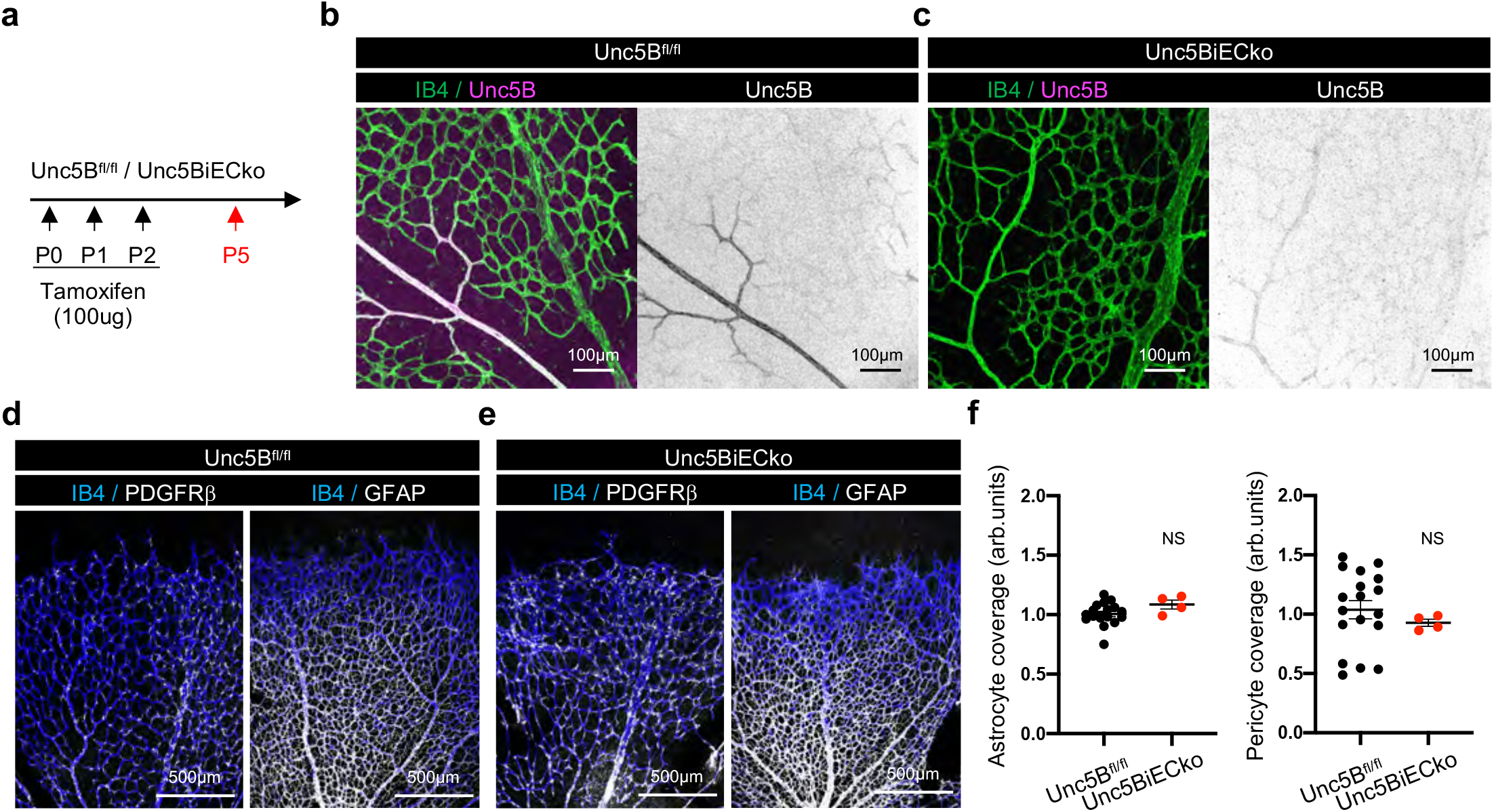
(a) *Unc5B* gene deletion strategy using tamoxifen injection in P5 neonates. (b-e) Immunofluorescence staining and confocal imaging on whole-mount P5 retinas with the indicated antibodies. (f) Quantification of GFAP+ astrocytes and PDGFRβ+ pericyte vascular coverage in P5 retinas. All data are shown as mean+/-SEM. Mann-Whitney U test was performed for statistical analysis between two groups.

**Supplemental Figure 3:**
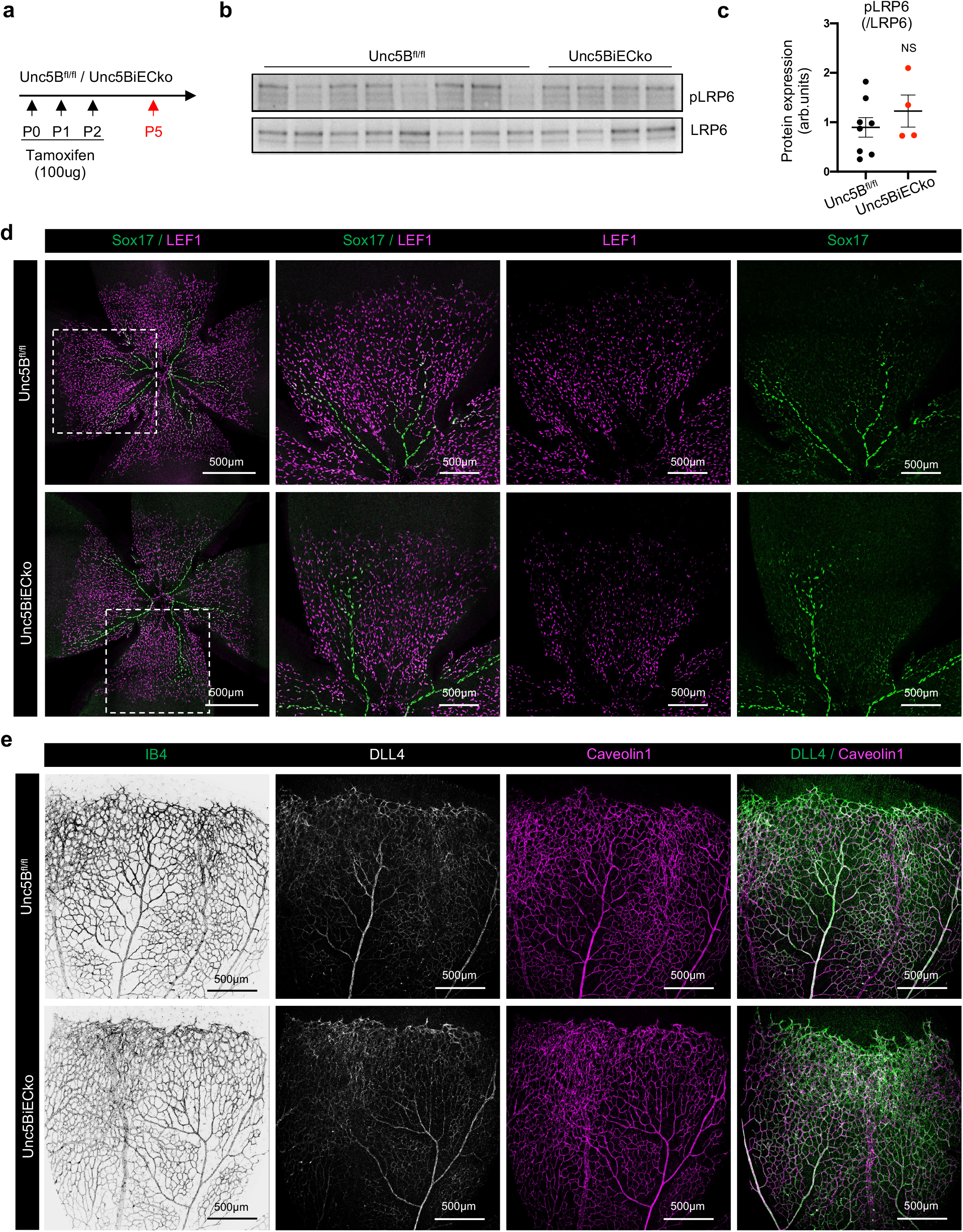
(a) *Unc5B* gene deletion strategy using tamoxifen injection in P12 neonates. (b,c) Western blot (b) and quantification (c) of P5 retina protein extracts. (d,e) Immunofluorescence staining and confocal imaging on whole-mount P5 retinas with the indicated antibodies. All data are shown as mean+/-SEM. Mann-Whitney U test was performed for statistical analysis between two groups.

**Supplemental Figure 4:**
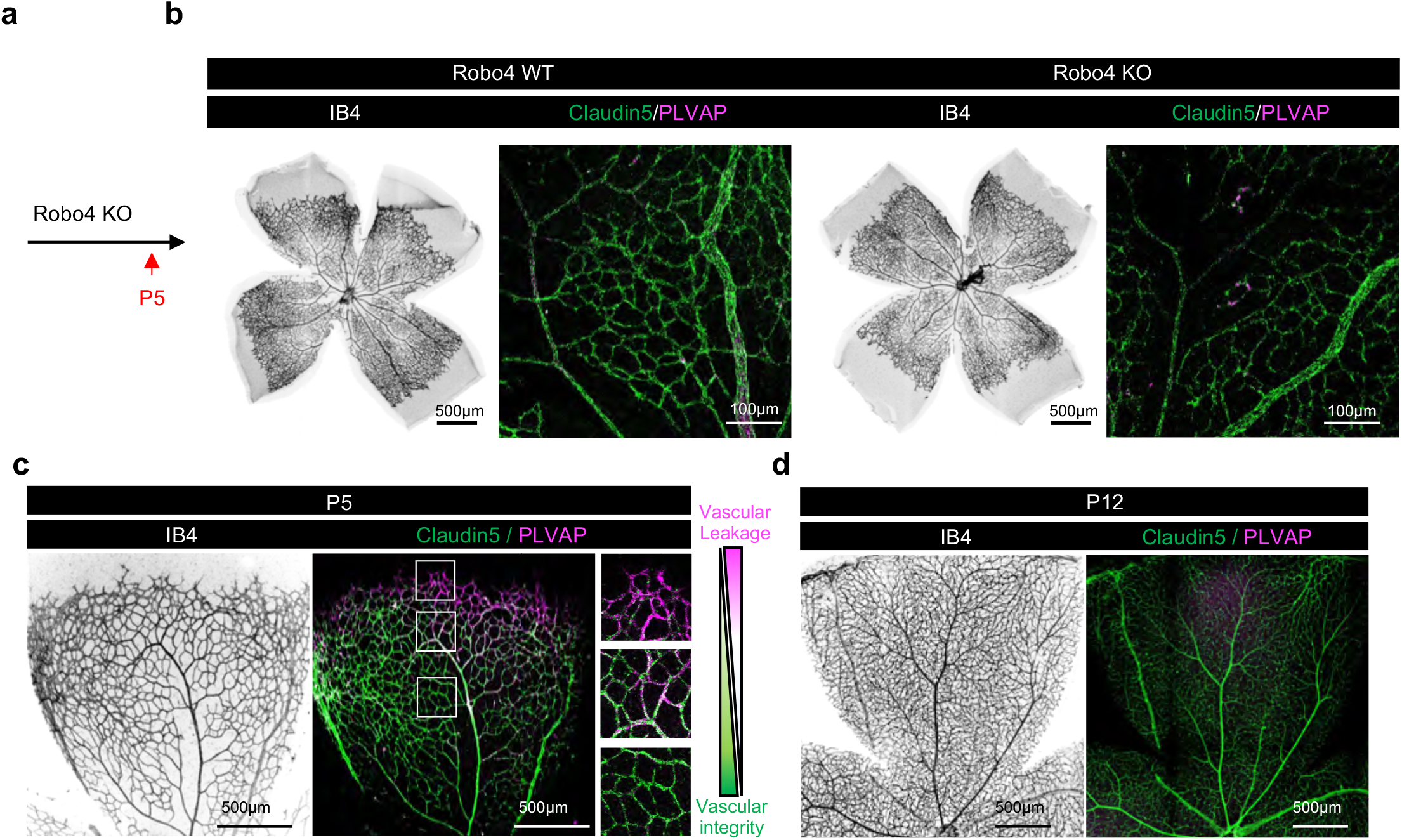
(a) *Robo4* gene deletion strategy in P5 neonates. (b-d) Immunofluorescence staining and confocal imaging on whole-mount P5 retinas with the indicated antibodies.

**Supplemental Figure 5:**
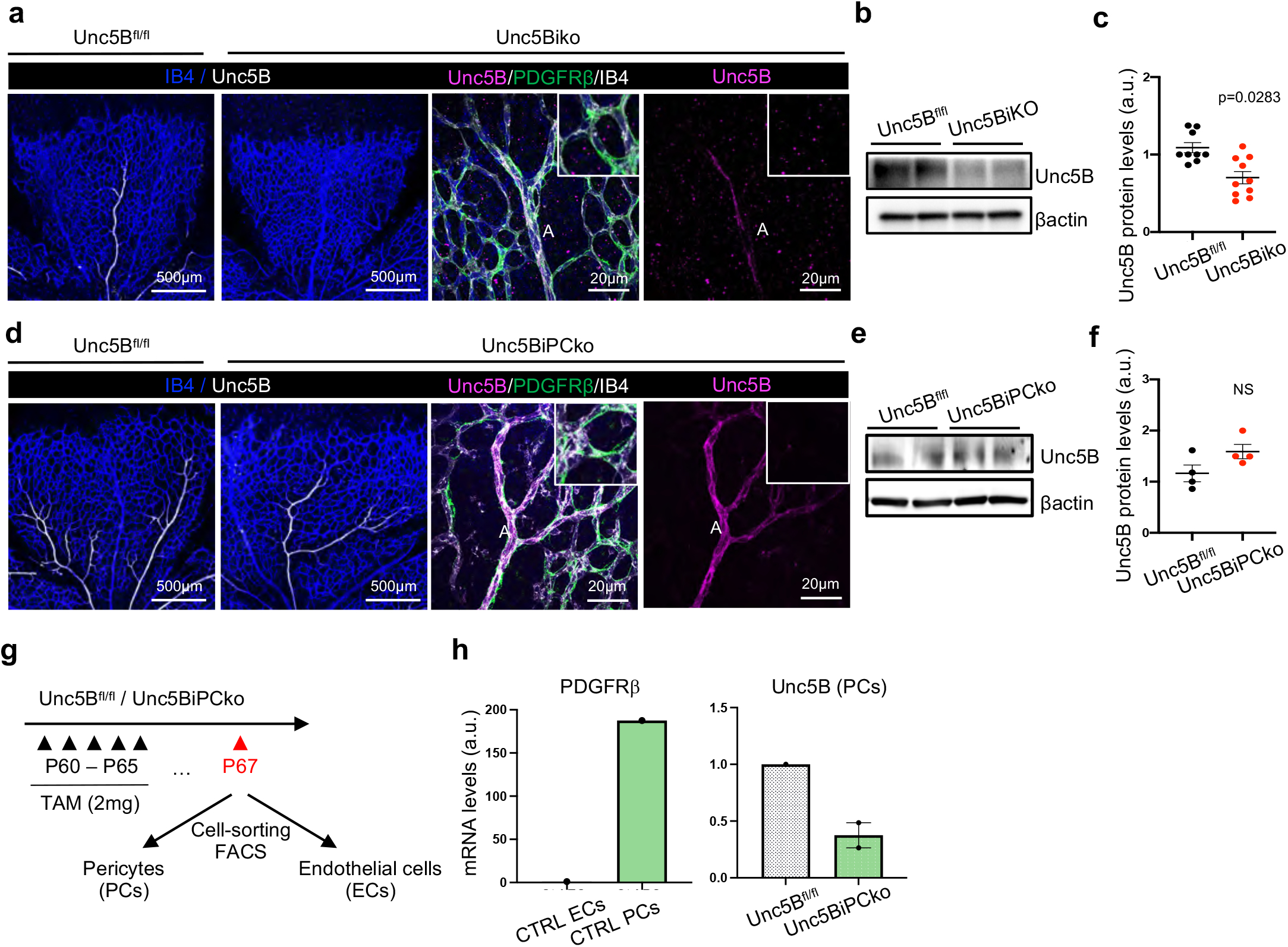
(a) Immunofluorescence staining and confocal imaging on whole-mount P5 retinas with the indicated antibodies. (b,c) Western blot (b) and quantification (c) of P5 retina protein extracts. (d) Immunofluorescence staining and confocal imaging on whole-mount P5 retinas with the indicated antibodies. (e,f) Western blot (e) and quantification (f) of P5 retina protein extracts. (g) Pericytes and endothelial cells isolation from CTRL or Unc5BiPCko brains. (h) qPCR analysis of mRNA extracted from ECs or pericytes isolated from CTRL or Unc5BiPCko brains. All data are shown as mean+/-SEM. Two-sided Mann-Whitney U test was performed for statistical analysis between two groups.

**Supplemental Figure 6:**
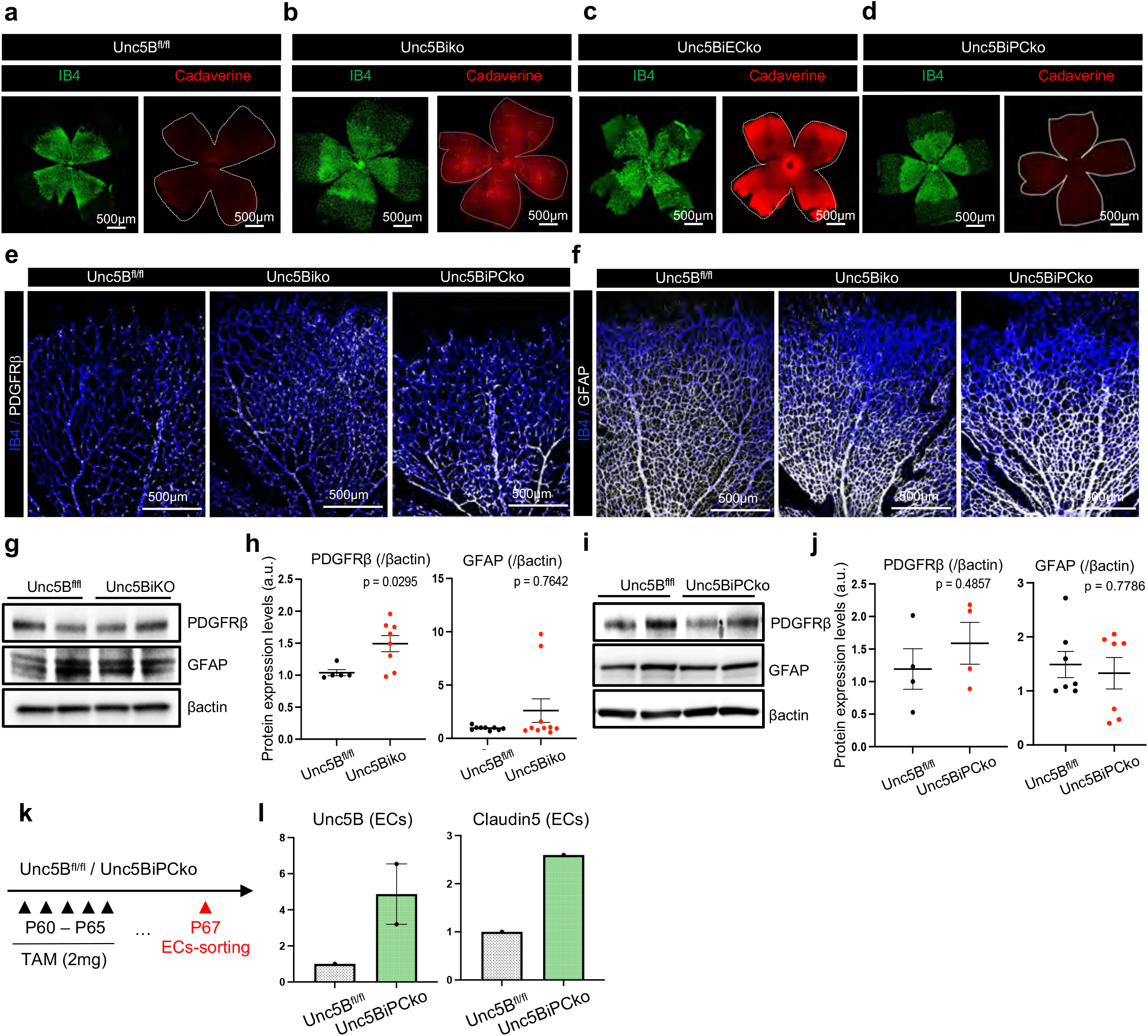
(a-d) Immunofluorescence staining and confocal imaging on whole-mount P5 retinas, 2h after i.p. cadaverine injection. (e,f) Immunofluorescence staining and confocal imaging on whole-mount P5 retinas with the indicated antibodies. (g,h) Western blot (g) and quantification (h) of P5 retina protein extracts. (i,j) Western blot (i) and quantification (j) of P5 retina protein extracts. (k) Pericytes and endothelial cells isolation from CTRL or Unc5BiPCko brains. (l) qPCR analysis of mRNA extracted from ECs or pericytes isolated from CTRL or Unc5BiPCko brains. All data are shown as mean+/-SEM. Two-sided Mann-Whitney U test was performed for statistical analysis between two groups. ANOVA followed by Bonferroni’s multiple comparisons test was performed for statistical analysis between multiple groups.

**Supplemental Figure 7:**
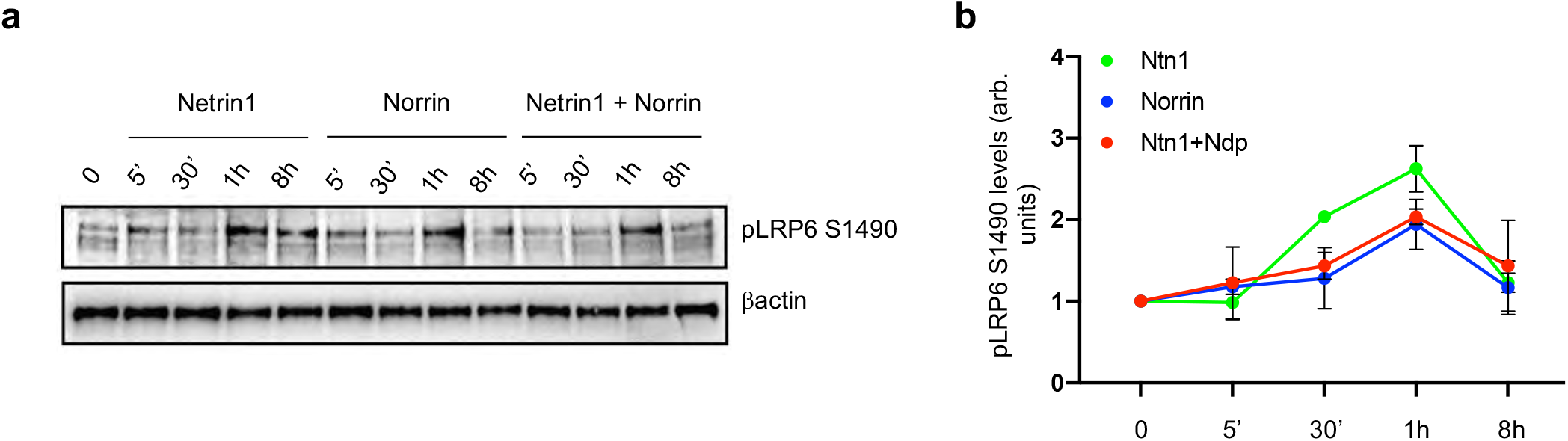
(a,b) Western blot (a) and quantification (b) of mouse brain ECs protein extract treated with recombinant mouse Netrin-1 protein or recombinant mouse Norrin protein, or a combination of both.

## ACKNOWLEDGEMENTS

This project has received funding from the NIH (1R01HL149343-01 to AE,), the Leducq Foundation (TNE ATTRACT, AE, LCW) and the European Research Council (ERC) (grant agreement No. 834161 to AE). KB was supported by a fellowship from the AHA (18POST34070109). JF was supported by a PhD fellowship from the AHA (Award number: 830299). We thank Pr. Patrick Mehlen and Dr. Nicolas Rama for sharing the *loxStoplox;hNtn1* mouse model.

## NOVELTY AND SIGNIFICANCE

### What is known

- The blood-retina barrier (BRB) tightly regulates transport across blood vessels, to protect retinal phototransduction.
- BRB integrity is regulated by the Norrin/β-catenin signaling pathway.
- Endothelial Netrin-1-Unc5B signaling regulates blood brain barrier (BBB) integrity via Wnt/β-catenin signaling.

### What new information does this article contribute

- Netrin-1 binding to endothelial Unc5B control BRB permeability via Norrin/β-catenin signaling.
- Mural cell Unc5B contributes to BRB regulation.
- Netrin-1-Unc5B signaling regulates Norrin/β-catenin mediated permeability via the Discs large homologue 1 (Dlg1) intracellular scaffolding protein.
- Increased Netrin1 levels induces conversion of leaky ECs Claudin-5-/PLVAP+ leaky state to a Claudin-5+/PLVAP-phenotype and increased BRB properties.

Impermeability of the functional blood retina barrier (BRB) impedes drug delivery for treatment in diseases, while loss of BRB permeability results in the development of retinal diseases like diabetic retinopathy and inflammatory disease. There is clinical need to develop therapeutic strategies that can increase or decrease BRB permeability. Using genetic mouse models and a blocking antibody, we show that Netrin-1 mediated Unc5B signaling regulates the BRB. The Norrin/β-catenin is the major pathway known to regulate BRB permeability. We found that disrupting Netrin-1 – Unc5B signaling via genetically inducible *Unc5B* and *Ntn1* deletion models, and antibody-mediated blockade of Netrin-1 binding to Unc5B decreased phosphorylation of the Norrin receptor LRP5 resulting in leakage of injected low molecular weight tracers and conversion of retina endothelial cells from a Claudin-5+/PLVAP-barrier-competent state to a Claudin-5-/PLVAP+ leaky phenotype. Netrin-1 – Unc5B regulated pLRP5 via the Discs large homologue 1 (Dlg1) intracellular scaffolding protein. Overexpressing β-catenin in *Unc5B* mutants rescued vascular leak and overexpressing *Ntn1* prompted barrier tightening. Pericyte Unc5B expression contributed to BRB permeability via endothelial Unc5B regulation. The data reveal endothelial Unc5B signaling as a CNS-wide endothelial barrier permeability control point that can be manipulated to decrease or increase barrier properties on demand.

